# Circulating tumor cell characterization and classification by novel combinatorial dual-color (CoDuCo) *in situ* hybridization and supervised machine learning

**DOI:** 10.1101/2024.05.08.592946

**Authors:** Lilli Bonstingl, Margret Zinnegger, Katja Sallinger, Karin Pankratz, Elisabeth Pritz, Corinna Odar, Christina Skofler, Christine Ulz, Lisa Oberauner-Wappis, Anatol Borrás-Cherrier, Višnja Somođi, Ellen Heitzer, Thomas Kroneis, Thomas Bauernhofer, Amin El-Heliebi

## Abstract

Metastatic prostate cancer is a highly heterogeneous and dynamic disease and practicable tools for patient stratification and resistance monitoring are urgently needed. Liquid biopsy analysis of circulating tumor DNA and circulating tumor cells (CTCs) are promising, but due to the diversity of resistance mechanisms, comprehensive testing is essential. Previously, we demonstrated that CTCs can be characterized by mRNA-based *in situ* padlock probe hybridization. Now, we have developed a novel combinatorial dual-color (CoDuCo) approach with increased multiplex capacity of up to 15 distinct markers, complemented by semi-automated image analysis and machine learning-assisted CTC classification. Here, we present three exemplary cases of patient samples in which the CoDuCo assay visualized diverse resistance mechanisms (AR-V7, neuroendocrine differentiation (SYP, CHGA, NCAM1)), as well as druggable targets and predictive markers (PSMA, DLL3, SLFN11). The combination of high multiplex capacity and microscopy-based single-cell analysis is a unique and powerful feature of the CoDuCo *in situ* assay. This synergy enables the identification and characterization of CTCs with epithelial, epithelial-mesenchymal, and neuroendocrine phenotypes, the detection of CTC clusters, and the visualization of CTC heterogeneity. In conclusion, the assay is a promising tool for monitoring the dynamic molecular changes associated with drug response and resistance in prostate cancer.

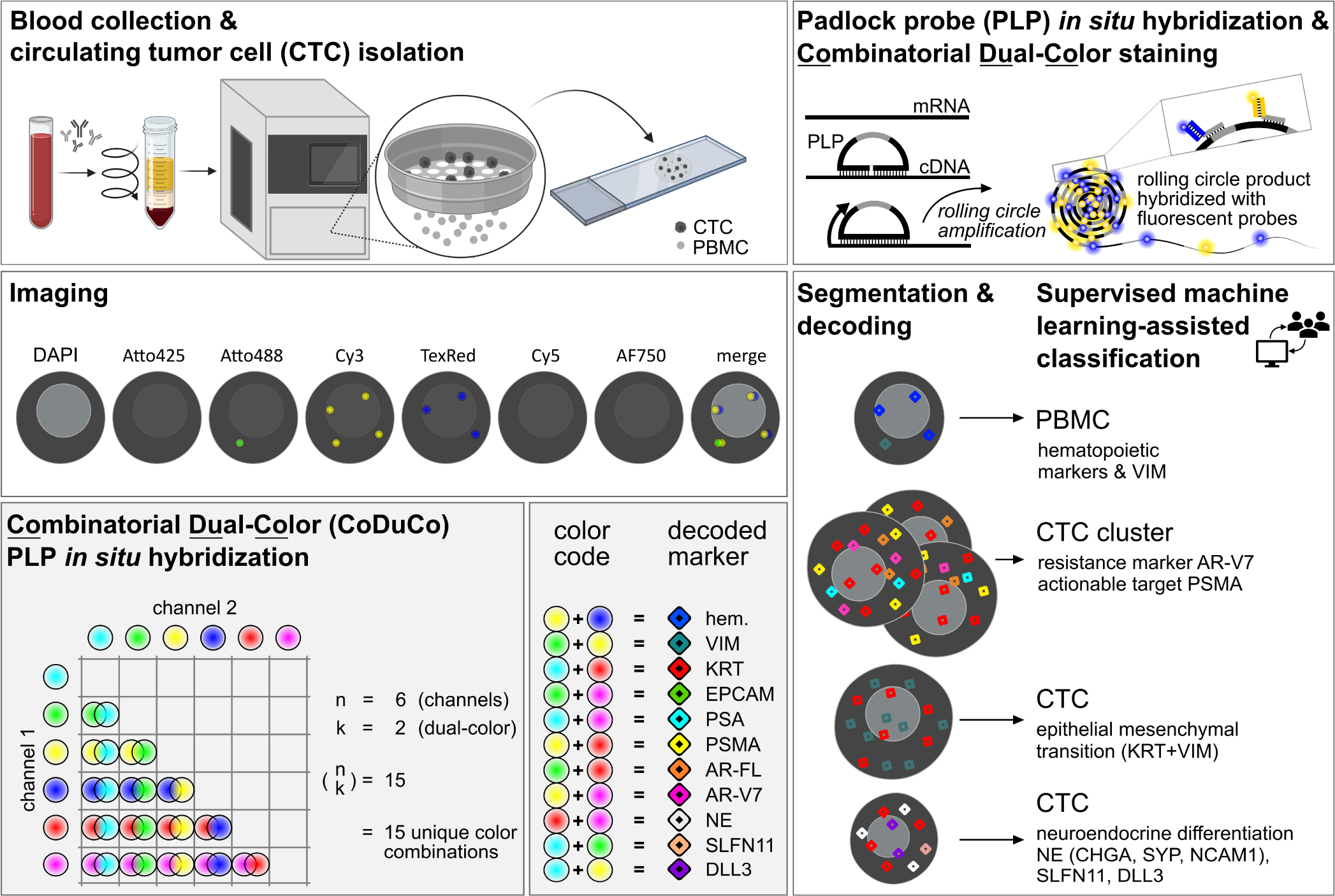

## INTRODUCTION

Prostate cancer (PC) is the third most frequently diagnosed solid cancer worldwide, with an estimated 1.4 million new cases reported in 2020 (1, 2). The risk for developing invasive PC increases with age and is highest for men older than 69 years (3, 4). With a world population that is growing and aging persistently, further increase of PC incidence is to be expected. Indeed, projections based on demographic changes and rising life expectancy suggest that annual new PC cases will increase to 2.9 million by 2040 (5).

Despite continuous improvements in treatment options for PC, the therapy of advanced PC is challenging, since the selective pressure created by new treatments also promotes the emergence of resistance mechanisms (6–8). Dysregulation of the androgen receptor (AR) pathway plays a key role in the development and progression of PC. Consequently, the most common treatment for advanced disease is androgen deprivation therapy (ADT). Although promising results are obtained, patients develop a high rate of resistance to ADT, hence classified as castration resistant prostate cancer (CRPC) (7). Novel AR-targeted treatments such as enzalutamide and abiraterone in combination with ADT represent effective therapy options for hormone-sensitive PC and CPRC patients (9). However, almost all patients acquire secondary resistance to novel AR-targeted treatments, leading to the progression of the fatal disease (9).

Resistance is frequently driven by aberrations of the AR signaling pathway, including *AR* gene amplification, mutations, and the expression of AR splice variants, particularly the splice variant AR-V7 (10–12). Additionally, AR-independent mechanisms like neuroendocrine transdifferentiation are on the rise as well (8).

To date, there are no coherent biomarker-based recommendations for optimal individualized treatment combinations and sequences of therapy lines, and clinically practicable tools for patient stratification and monitoring of drug resistance are unavailable (13, 14). There is an urgent need to investigate a multitude of resistance mechanisms, which could be exploited as predictive biomarkers. For example, AR-V7-positive patients are more likely to benefit from taxane-based chemotherapy than AR-targeted drugs, while platinum-based chemotherapy may be indicated in patients with neuroendocrine PC (15, 16). Moreover, other treatments, such as drugs targeting PSMA or DLL3-expressing tumor cells, are increasingly available or under investigation (17, 18).

In recent years, the concept of liquid biopsy gained tremendous attention as a minimally invasive way to monitor disease state using different analytes such as circulating tumor cells (CTCs) and circulating tumor DNA (ctDNA) (19–21). However, a major challenge in liquid biopsies lies in simultaneously investigating a broad spectrum of resistance mechanisms and predictive biomarkers in PC. Relevant alterations range from genetic aberrations (e.g. *AR*-gain or mutations) to transcriptional changes such as alternative splicing (e.g. AR-V7), expression of PSMA and SLFN11, or upregulation of neuroendocrine markers (e.g. SYP, CHGA, NCAM1, and DLL3) in cancers cells. Therefore, a comprehensive liquid biopsy approach is needed to capture various druggable targets and resistance mechanisms. While ctDNA analysis excels in uncovering genetic alterations, transcriptional regulation analysis of single gene loci by ctDNA nucleosome patterns remains challenging (22). Previously, we have demonstrated that resistance mechanisms in CTCs can be characterized by mRNA-based *in situ* padlock probe (PLP) hybridization, using an assay that provided broad expression data of AR, AR-V7, and PSA in CTCs of PC patients (23). The main technical challenges of *in situ* PLP-based CTC analysis were time-intensive manual evaluation of *in situ* data and restricted multiplexability of the existing technique (e.g. limitation to detect 3 transcripts on a 4-channel fluorescence microscope).

Building on this, we set out to provide an advanced multiplex mRNA-based *in situ* assay that reveals additional predictive biomarkers in CTCs. Through machine learning algorithms, the time-intensive image evaluation of CTCs is streamlined, thereby overcoming the aforementioned limitations.

Here, we present our achievements in developing a multiplex mRNA-based CTC assay examining multiple biomarkers with significant predictive power, including PSMA, PSA, AR, AR-V7, and neuroendocrine markers SYP, CHGA, and NCAM1. Our innovative approach introduces a novel combinatorial dual-color (CoDuCo) *in situ* hybridization assay with increased multiplex capacity of up to 15 targets, semi-automated image analysis, and machine learning-assisted CTC classification.

## MATERIALS AND METHODS

### Patient Sampling and Ethics

The study enrolled patients with advanced metastatic PC at the Division of Oncology, Department of Internal Medicine, Medical University of Graz (Austria), following the principles of the World Medical Association Declaration of Helsinki. The study was approved by the ethics committee (EK 31-353 ex 18/19) and written informed consent was obtained from all patients and healthy controls. To avoid contamination by epithelial cells, one extra blood tube with 2.5 ml of blood was collected for non-cell-based analyses before collecting the blood samples for cell-based analyses. For isolation of peripheral blood mononuclear cells (PBMCs) from healthy controls, blood samples were collected in VACUETTE blood collection tubes K3E K3EDTA (9 mL blood) (Greiner Bio-One, Kremsmünster, Austria). For CTC enrichment, blood samples were collected in 8.5 ml BD Vacutainer ACD-A tubes (BD Switzerland Sarl, Eysins, Switzerland). All blood samples were collected following the CEN/TS 17390-3 standards to ensure defined pre-analytical parameters as described previously (24).

### Cell line and PBMC sample preparation

PC cell lines VCaP (kindly provided by Martina Auer, Medical University of Graz, Graz, Austria) and PC-3 (American Type Culture Collection (ATCC), Manassas, VA, USA) were cultured as described in detail by Hofmann, Kroneis and El-Heliebi (25). The lung cancer cell line NCI-H1299 (kindly provided by Eva Obermayr, Medical University of Vienna, Vienna, Austria) was cultured in RPMI (Roswell Park Memorial Institute) 1640 Medium (Thermo Fisher Scientific, Waltham, MA, USA) with 10% FBS and 1% Penicillin/Streptomycin at 37°C and 5% CO_2_. All cell lines were harvested as published previously (25).

For *in situ* assay validation experiments, the PBMC fraction of healthy donors’ blood samples in VACUETTE blood collection tubes K3E K3EDTA was isolated by density gradient centrifugation as described previously (23). PBMCs, VCaP, PC-3, and NCI-H129 cells were fixed in 3.7% formaldehyde (Sigma-Aldrich, St Louis, MO, USA, catalog number F1635) in PBS (Thermo Fisher Scientific, catalog number 10010015) for 5 min, resuspended in 1×PBS, and 1×10^5^ cells were transferred to SuperFrost Plus microscope slides (Thermo Fisher Scientific, catalog number J1800AMNT) by cytocentrifugation using a Hettich Universal 32 benchtop centrifuge. Slides were dried over night at room temperature and stored at −80°C.

### CTC enrichment and sample preparation for *in situ* analysis

CTCs were enriched from 7.5 ml of blood samples collected in ACD-A tubes using the Cytogen Smart Biopsy Cell Isolator (Cytogen Inc., Seoul, Korea) following the manufacturer’s protocol. In short, the double negative selection workflow involves incubation with a leukocyte and erythrocyte depletion cocktail and subsequent density gradient centrifugation. Samples are then forwarded to the automated Smart Biopsy Cell Isolator, which performs a size-based filtration using a high-density microporous (HDM) chip, retrieval of the enriched cell fraction from the HDM chip, and automated transfer to a reaction tube (26). The cells were fixed with 2% formaldehyde for 5 minutes and centrifuged on Micro Slide Glass Frontier FRC-01 (Matsunami Glass Industry Ltd, Osaka, Japan) using a Shandon Cytospin 2. The cells were washed 3 times with PBS, dried over night at room temperature and stored at −80°C. This procedure was applied to patient blood samples, healthy controls’ blood samples, and healthy controls’ blood samples spiked with 500 VCaP and 500 PC-3 cells.

### CoDuCo *in situ* PLP hybridization

*In situ* PLP hybridization with a novel CoDuCo staining approach was used to visualize transcripts in cells. *In situ* PLP hybridization starts with targeted reverse transcription. Then, PLPs are hybridized to the cDNA and ligated to form closed circular molecules. The PLP sequence is amplified by rolling circle amplification and finally, bridge probes and fluorescently labelled readout detection probes are hybridized to the resulting rolling circle products (RCPs) (25, 27, 28).

#### Selection of genes

CoDuCo *in situ* hybridization was used to visualize hematopoietic transcripts *PTPRC* (protein tyrosine phosphatase receptor type C (CD45), GenBank accession number NM_002838.5), *ITGAM* (integrin subunit alpha M (CD11B), NM_000632.4), *FCGR3A* (Fc gamma receptor IIIa (CD16), NM_001127596.2), *FCGR3B* (Fc gamma receptor IIIb (CD16), NM_001271035.2), *CD4* (CD4 molecule, NM_001195017.3), and *ITGB2* (integrin subunit beta 2 (CD18), NM_000211.5); epithelial transcripts *EPCAM* (epithelial cell adhesion molecule, NM_002354.3), *KRT8* (keratin 8, NM_001256293.2), *KRT18* (keratin 18, NM_000224.3), and *KRT19* (keratin 19, NM_002276.5); prostate-specific transcripts *KLK3* (kallikrein related peptidase 3 (prostate specific antigen PSA), NM_001030047), *FOLH1* (folate hydrolase 1 (prostate specific membrane antigen PSMA), NM_004476.3), *AR-FL* (androgen receptor full length, NM_000044.3), and *AR-V7* (androgen receptor splice variant 7, FJ235916.1); neuroendocrine transcripts *SYP* (synaptophysin, NM_003179.3), *CHGA* (chromogranin A, NM_001275.4), *NCAM1* (neural cell adhesion molecule 1, NM_001242607.2), and *DLL3* (delta like canonical Notch ligand 3, NM_203486.3). In addition, *VIM* (vimentin, NM_003380.5) and *SLFN11* (schlafen family member 11, NM_001376010.1) were visualized.

#### Probe design

Reverse transcription primers with a length of 15-25 nucleotides were designed using CLC Main Workbench software (CLC Bio workbench version 7.6, QIAGEN, Hilden, Germany) based on the guidelines published by Weibrecht et al. (29). Primer binding sites close to the PLP binding sites were preferred and an overlap of up to 6 nucleotides was allowed. Up to 6 locked nucleic acids (LNA)-modified nucleotides were included in selected primers to increase binding strength. No overlap of LNA-modified nucleotides with the PLP-binding site was allowed.

PLPs consist of 3’ and 5’ target-binding arms linked by a central backbone. PLPs were designed using a Python software package developed by the Mats Nilsson Lab, Stockholm University (https://github.com/Moldia/multi_padlock_design), with an arm length of 15 nucleotides and melting temperature between 65°C and 75°C (28). Some PLP binding sites were determined manually, using CLC Main Workbench software based on the guidelines published by Weibrecht et al. (29). Manually designed PLPs had 15-19 nucleotide long target-binding arms, covering a binding site of 30-38 nucleotides in total. The PLP backbones contain a 17-20 nucleotide ID sequence unique for each gene. To increase sensitivity of the assay, each gene was targeted by up to 44 reverse transcription primers and up to 20 PLPs.

Bridge probes were used for indirect hybridization of 5’ fluorescently labelled readout detection probes to the RCP. They consist of the 17 nucleotide ID sequence, which binds to the RCP, a 2-nucleotide linker, and the reverse complementary 20 nucleotide sequence of the respective readout detection probes.

Oligonucleotides were ordered from IDT (Integrated DNA Technologies, Coralville, IA, USA) and stored at −20°C as 100 µM stocks or 10 µM dilutions in nuclease-free water (Thermo Fisher Scientific, catalog number AM9930) or IDTE buffer pH 8 (Integrated DNA Technologies). PLPs were ordered 5’ phosphorylated.

#### Buffers for in situ hybridization

Diethyl pyrocarbonate (DEPC, Sigma-Aldrich, catalog number D5758) was used to remove RNase activity in ultrapure water (H_2_O). DEPC-H_2_O (1 ml DEPC in 1 l H_2_O) was incubated overnight at room temperature and then autoclaved to deactivate DEPC. DEPC-H_2_O was used to prepare DEPC-PBS (phosphate buffered saline) and DEPC-PBS-Tween (0.05% Tween-20, Sigma-Aldrich, catalog number 822184).

#### In situ hybridization and imaging

Prefixed slides were thawed for 3 minutes, fixed again with 3.7% formaldehyde in 1×DEPC-PBS for 15 minutes, and washed with 1×DEPC-PBS-Tween for 2 minutes. Slides were then dehydrated through ascending ethanol series (70%, 85%, and 100% ethanol in DEPC-H_2_O for 2 minutes each). 50 µl SecureSeal hybridization chambers (Thermo Fisher Scientific, catalog number S24732) were mounted on completely air-dried slides to cover the cytospinned cells. Cells were rehydrated with 1×DEPC-PBS-Tween for 5 minutes, permeabilized with 0.1 M HCl (hydrochloric acid, Merck Chemicals and Life Sciences, Vienna, Austria, catalog number 1.09970.0001) in DEPC-H_2_O for 5 minutes and washed twice with 1×DEPC-PBS-Tween for 5 minutes each.

The reaction mix for reverse transcription (RT) contained 40 U/μl TranscriptMe Reverse Transcriptase (DNA-Gdansk, Gdansk, Poland, catalog number RT32-010), 2 U/μl RiboLock RNase inhibitor (Thermo Fisher Scientific, catalog number EO0381), 0.5 mM dNTPs (Sigma-Aldrich, catalog number D7295), 0.1 μM of each reverse transcription primer and 0.4 μg/μl BSA (Thermo Fisher Scientific, catalog number B14) in RT buffer (DNA-Gdansk). Reverse transcription was carried out at 45°C for 3 hours, followed by fixation with 3.7% formaldehyde in 1×DEPC-PBS for 10 minutes, and two washing steps with 1×DEPC-PBS-Tween for 2 minutes each.

The reaction mix for PLP hybridization and ligation contained 1 U/μl Ampligase (Biozym Biotech Trading, Vienna, Austria, catalog number 111075), 0.8 U/μl RNase H (Thermo Fisher Scientific, catalog number EN0201), 0.4 μg/μl BSA, 0.1 μM of each PLP, 0.05 M KCl (potassium chloride, Sigma-Aldrich, catalog number P9333) and 20% formamide (Sigma-Aldrich, catalog number F9037) in Ampligase buffer. Incubation was started at 37°C for 30 minutes for RNA digestion, followed by 45 minutes at 45°C for PLP hybridization and ligation. Samples were washed once with pre-warmed 2×SSC-Tween (saline sodium citrate buffer, Thermo Fisher Scientific, catalog number 15557044; 0.05% Tween-20) at 37°C for 5 minutes and twice with 1×DEPC-PBS-Tween at room temperature for 2 minutes each.

The reaction mix for rolling circle amplification contained 2 U/μl φ29 Polymerase (Thermo Fisher Scientific, catalog number EP0091), 0.4 μg/μl BSA, 0.25 mM dNTPs, and 5% Glycerol (Carl Roth, Karlsruhe, Germany, catalog number 3783.1) in φ29 Polymerase buffer. Amplification was performed overnight at room temperature, followed by two washes with 1×DEPC-PBS-Tween for 2 minutes each.

Bridge probes were hybridized at a final concentration of 0.1 µM in a hybridization buffer of 20% formamide in 2×SSC at 37°C for 60 minutes, followed by two washes with 2×SSC for 2 minutes each. Readout detection probes were hybridized at a final concentration of 0.1 µM together with 2 µg/ml DAPI (Thermo Fisher Scientific, catalog number D21490) in a hybridization buffer of 20% formamide in 2×SSC at 37°C for 30 minutes, followed by two washes with 1×DEPC-PBS-Tween for 2 minutes each.

SecureSeal hybridization chambers were removed from the slides and the cytospins were covered with SlowFade Gold Antifade Mountant (Thermo Fisher Scientific, catalog number S36936) and a coverslip for imaging.

To increase signal to noise ratios we implemented a mathematical subtraction of background fluorescence. To do so, we first imaged the slides, then removed *in situ* signals by formamide stripping, rescanned the remaining background fluorescence and subtracted this background from the originally scanned images. In detail, slides were soaked in 1×DEPC-PBS to gently remove coverglass and mounting medium, dehydrated through ascending ethanol series (70%, 85%, and 100% ethanol in DEPC-H_2_O for 2 minutes each) and air-dried. 50 µl SecureSeal hybridization chambers were mounted on the slides. Samples were rehydrated with 1×DEPC-PBS-Tween for 5 minutes and incubated three times with prewarmed 100% formamide at 37°C, followed by two washes with 1×DEPC-PBS-Tween for 2 minutes each. To enhance the DAPI staining, samples were incubated with 2 µg/ml DAPI in 1×DEPC-PBS at 37°C for 30 minutes. Samples were washed twice with 1×DEPC-PBS-Tween for 2 minutes each. The SecureSeal hybridization chambers were removed, and the samples were mounted for imaging as described before.

Unless noted otherwise, all steps were performed at room temperature.

#### Imaging

Slides were imaged using Slideview VS200 digital slide scanners (Evident, Tokio, Japan). The scanners were equipped with external LED light sources Xcite Xylis or Xcite Novem (Excelitas, Mississauga, Canada), Olympus universal-plan extended apochromat 40x objectives (UPLXAPO40x, 0.95 NA/air; Olympus), and Hamamatsu ORCA-Fusion digital sCMOS cameras (C14440-20UP, 2304 × 2304 (5.3 Megapixels), 16 bit; Hamamatsu City, Japan). DAPI, Cy5, and AF750 were imaged using a Semrock penta filter (AHF-LED-DFC3C5C7-5-SBM; AHF, Tübingen-Pfrondorf, Germany / Semrock / IDEX Health & Science, Oak Harbor, Washington, USA) with excitation wavelengths of 378/52 nm, 635/18 nm, and 735/28 nm, and emission wavelengths of 432/36 nm, 685/42 nm, and 809/81 nm. Atto425, Atto488, Cy3, and TexasRed were imaged using Spectrasplit filters (Kromnigon, Goteborg, Sweden) with excitation wavelengths of 438/24 nm, 509/22 nm, 550/10 nm, and 578/21 nm, and emission wavelengths of 482/25 nm, 544/24 nm, 565/16 nm, and 641/75 nm. Exposure times were adjusted depending on the used light source (DAPI 0.8-3 ms, Atto425 50-70 ms, Atto488 50-70 ms, Cy3 30-70 ms, TexasRed 40-70 ms, Cy5 20-200 ms, AF750 100-300 ms). Extended focus imaging with z-range of 7.41 µm and z-spacing of 0.84 µm (9 z-planes) was used to depict in-focus *in situ* signals in a single plane.

#### Image analysis

Original and background scan images were converted to TIF file format (LZW compression) in reduced resolution by 8×8 binning as well as in full resolution. The 8×8-binned DAPI image was segmented in CellProfiler version 4 (Broad Institute of MIT and Harvard, Cambridge, MA, USA) (30) to detect regions of interests (ROIs) occupied by nuclei. The coordinates of ROI bounding boxes were exported. In addition, a binary image of each ROI was cropped and saved.

The Python package pyStackReg was used for the registration of original and background images (31). During pre-alignment, a rigid body transformation matrix was computed for the 8×8 binned DAPI images. The matrix was multiplied by 8 for upscaling and applied to register the full resolution background image. ROIs were then cropped from the full resolution images based on the coordinates determined in CellProfiler. For optimal image alignment, the rigid body registration based on the DAPI channel was repeated for each cropped full resolution ROI image.

The cropped full resolution images and the binary ROI images were then subjected to a second CellProfiler pipeline for cell segmentation, *in situ* signal detection and decoding, and cell classification. First, for all channels except DAPI, the background images were subtracted from the original images to decrease autofluorescence of cytoplasm and other objects such as erythrocytes, thereby increasing the relative intensity of *in situ* signals (32). The binary ROI image was used to mask the DAPI image to remove nuclei of neighboring ROIs with overlapping bounding boxes. High and low intensity nuclei were identified by separate instances of the “IdentifyPrimaryObjects” module and were then combined into a single object set. Cell borders were identified by expanding nuclei by 12 px (2 µm) or 32 px (5 µm) depending on nucleus area below or above 2800 px (73.6 µm^2^; radius 4.8 µm). In samples that contained cells with particularly large cytoplasmic volume, an additional propagation step, guided by autofluorescence in the Atto488 background scan, was included. *In situ* signals were identified using adaptive minimum cross-entropy thresholding. *In situ* signals were shrunk to a uniform diameter of 3 px to 5 px with local intensity maxima at the object center. For *in situ* signal decoding, the “Relate Objects” module was deployed for each utilized dual-color code to identify colocalized signals. In areas with high density of *in situ* signals, overlapping transcript-spots can lead to decoding problems and spurious calling of multiple transcripts. To minimize this effect, transcripts with expected high expression were used to mask potential false-positive transcripts in a multi-level process. VIM and KRT were used to mask AR-FL, i.e. AR-FL signals that overlapped with VIM and/or KRT were removed. VIM, KRT, and AR-FL were used to mask PSA. VIM, KRT, AR-FL, and PSA were used to mask EPCAM, PSMA, AR-V7, pooled neuroendocrine markers NE, SLFN11, and DLL3. The “RelateObjects” module was also used to assign the decoded *in situ* signals to appointed cells. Three sources of false-positive *in situ* signals were observed. These sources were identified by additional CellProfiler modules and used to mask potentially false-positive *in situ* signals. First, objects with high autofluorescence in multiple channels were identified if detectable in both the Cy5 and TexasRed background scan. Second, colocalized *in situ* signals that were detectable in >4 channels, were defined as unspecific signals. Third, empty image areas at the borders of the background scans resulting from image registration. For identification of nuclei, *in situ* signals, highly autofluorescent background objects, and, where applicable, propagated cell borders, the lower bounds on threshold and threshold correction factor were adjusted manually for each channel and sample. The *in situ* signal count for each cell was exported to a csv spreadsheet and the results were visualized using the “OverlayOutlines” module. When analyzing patient samples, cells were classified directly in CellProfiler based on their *in situ* signal counts, using the “ClassifyObjects” module and a model trained on control samples in CellProfiler Analyst version 3 (Broad Institute of MIT and Harvard) (33).

#### Machine learning-based classification

To create a training and test dataset for the CellProfiler Analyst Classifier tool, 6 blood samples of healthy controls without (n=3) or with (n=3) spiked-in VCaP and PC-3 cells were enriched for CTCs and transcripts were visualized by *in situ* hybridization. After image analysis in CellProfiler, the full dataset was annotated by expert evaluation. Cells were categorized in 5 classes: CTCs (*in situ* signals detected for epithelial and/or prostate-specific markers), PBMCs (*in situ* signals detected for hematopoietic markers), artefacts (objects without nucleus, e.g. dust particles), *in situ* false-positive cells (*in situ* signals were detected by CellProfiler, but were rated as unspecific by the expert, e.g. due to high autofluorescence), and *in situ* negative cells (no *in situ* signals detectable by CellProfiler). The dataset was split into a training and test dataset. For the training dataset, 500 PBMCs were randomly chosen from the samples without spiked-in tumor cells. 200 CTCs were chosen randomly from the samples with spiked-in tumor cells. For the remaining classes, 100 cells each were randomly chosen from all samples. Cells with over- or undersegmentation of nuclei, and cells with both true-positive and false-positive *in situ* signals were not included in the training dataset. The training dataset was used in CellProfiler Analyst to train a random forest classifier, based on the decoded *in situ* signals per cell. The classifier was then evaluated in the test dataset. The classifier model was saved and integrated in the CellProfiler pipelines for image analysis and classification of patient samples. The patient samples were subjected to expert revision to identify false-positive or false-negative CTCs and, where necessary, correct the RCP counts in CTCs.

### AdnaTest ProstateCancerSelect AR-V7

The AdnaTest ProstateCancerSelect AR-V7 (QIAGEN, Hilden, Germany) was used according to manufacturer’s guidelines as described previously (24). In short, 5 ml of whole blood collected in PAXgene Blood ccfDNA Tubes (PreAnalytiX, Hombrechtikon, Switzerland) were used for immunomagnetic enrichment of CTCs. mRNA was isolated from the lysate of pre-enriched CTCs, and cDNA (20 µl) created by reverse transcription. The cDNA was then subjected to preamplification PCR in triplicates (6.25 µl cDNA per reaction). Finally, qRT-PCR was performed to detect CD45, GAPDH, PSA, PSMA, AR, and AR-V7. Patient samples were considered positive for the respective marker if it was detected in at least one of the cDNA triplicates.

### Data analysis and visualization

A number of Python libraries, including numpy and pandas, were used for data analysis and visualization (34–36). Median and interquartile range (IQR) of CoDuCo *in situ* signals per cell (RCPs/cell) were calculated using the Python library pandas. Shapiro-Wilk test (scipy library) and Q-Q-plots (statsmodels library) were used to test data for normal distribution (37, 38). As the data were not normally distributed, Kruskal-Wallis test (scipy) was performed to find significant differences, followed by pairwise comparisons using Dunn’s test (scikit_posthocs library), with p-value adjustment for multiple comparisons using the Benjamini and Hochberg method (39) and the resulting q-values were reported, with q-values <0.05 considered statistically significant. To evaluate the performance of the random forest classifier for CTC detection, the sklearn library was used to calculate confusion matrix, multilabel confusion matrix, precision (positive predictive value), recall (sensitivity), f1-score, and support (40). Furthermore, specificity and Matthews Correlation Coefficient were calculated (41, 42). The Python libraries plotly, matplotlib, and seaborn were used to create plots and figures, which were finalized in Inkscape (43, 44). PyCharm and Jupyter were used for Python projects and virtualenv and conda for managing virtual environments (45).

## RESULTS

### Decoding of CoDuCo *in situ* signals

*In situ* PLP hybridization can be used to visualize transcripts in CTCs. Aiming to increase the number of detectable transcripts, we developed a novel staining approach. With a conventional approach, a seven-channel fluorescence microscope can detect a maximum of six mRNA markers along with DAPI-stained nuclei. To address this limitation, we introduced the CoDuCo approach, which employs a two-color code for *in situ* signal detection. Given six fluorescence channels for *in situ* signals (n=6) and a two-color code (k=2), the total number of distinct combinations is (*^n^*)*=15*.

We had noticed bleed-through of Cy5 into the TexasRed channel. To minimize the risk for decoding errors, we made a strategic choice and opted to use TexasRed exclusively for detecting hematopoietic markers in PBMCs, in combination with Cy3. This decision resulted in a total of 11 unique color combinations. **Figure 1** provides a summary of all markers and their corresponding color codes. Additionally, it illustrates how a tumor cell and a PBMC were identified based on the decoded *in situ* signals. To cover a broad range of PBMCs, we used a panel of five hematopoietic markers, that were all detected by the TexasRed+Cy3 combination, namely, PTPRC (CD45), ITGAM (CD11B), FCGR3A and FCGR3B (CD16), CD4, and ITGB2 (CD18). Similar to the pooled hematopoietic markers, we detected pooled KRT markers (KRT8, KRT18, and KRT19) using the Atto425+Cy5 code, and pooled neuroendocrine markers NE (SYP, CHGA, and NCAM1) using Cy5+AF750. The remaining color codes were used to detect VIM (Atto488+Cy3), EPCAM (Atto488+AF750), PSA (Atto425+AF750), PSMA (Cy3+Cy5), AR-FL (Atto488+Cy5), AR-V7 (Cy3+AF750), SLFN11 (Atto425+Atto488), and DLL3 (Atto425+Cy3).

**Figure 1.**
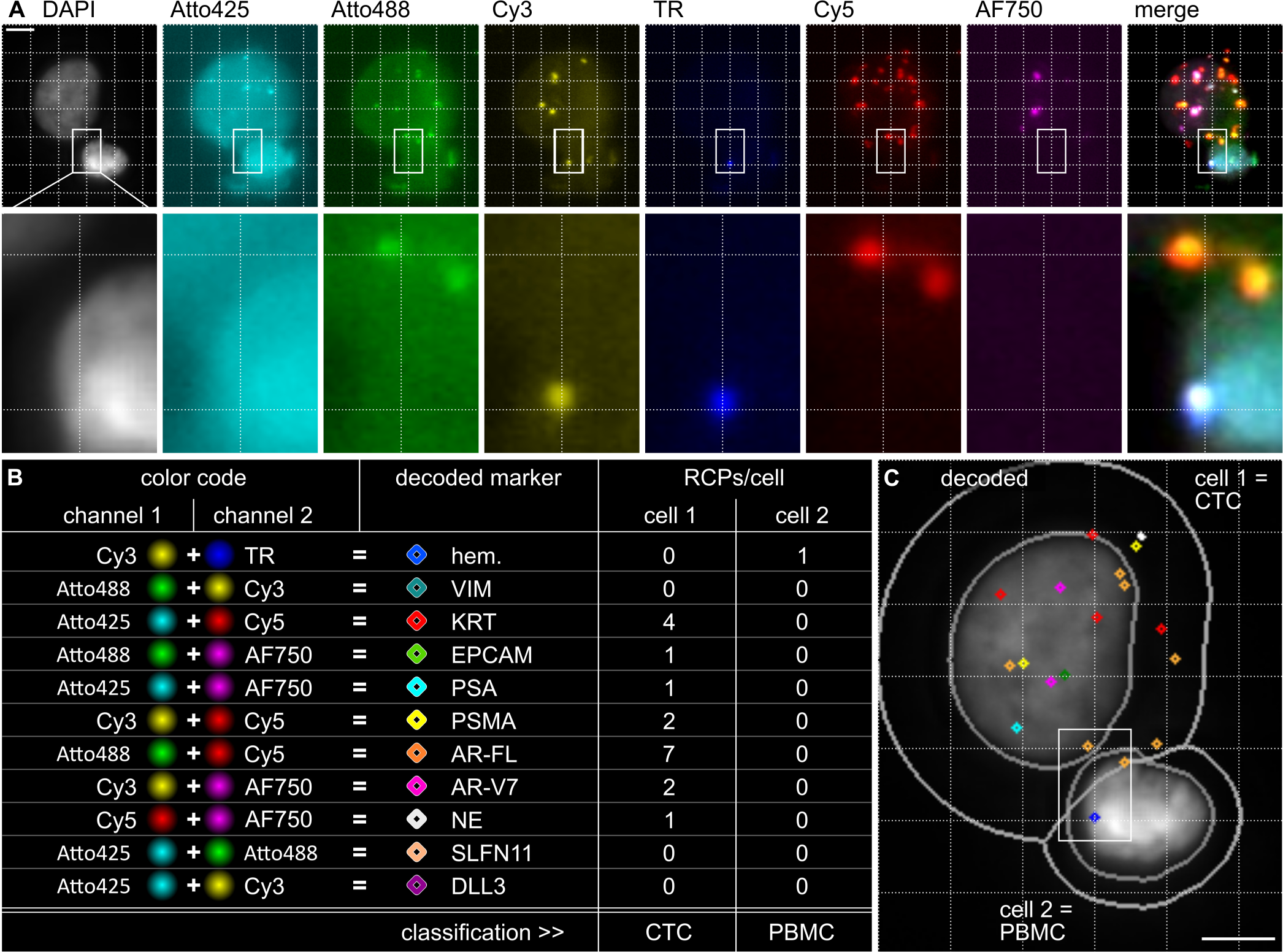
Decoding CoDuCo in situ signals to identify CTCs and PBMCs. (A) Pseudocolored images of two cells with DAPI-stained nuclei and CoDuCo in situ signals in 6 channels. The dotted 5 µm grid is a visual aid, guiding the eye to recognize colocalized in situ signals. One region is highlighted and shown in higher magnification. (B) Decoding scheme for all markers and their expression levels [RCPs/cell] in the two cells. E.g. in situ signals that are visible at the exact same x and y coordinates in the Cy3 and TR (TexasRed) channel, can be decoded as hematopoietic (hem.) markers. The respective colored diamond-shapes are used in (C) to visualize decoded in situ signals, together with gray outlines of nuclei and cell borders on the DAPI image. PBMCs can be identified based on their expression of hematopoietic markers, and CTCs can be identified based on the expression of epithelial (KRT and EPCAM) and/or prostate-specific markers (PSA, PSMA, AR-FL, AR-V7). Scale bar 5 µm.

### Validation of CoDuCo for CTC characterization

To validate the novel CoDuCo *in situ* assay, we applied it to healthy control PBMCs, PC cell lines VCaP and PC-3, and non-small cell lung cancer cell line NCI-H1299. We detected CoDuCo *in situ* signals in 89% of PBMCs (n=7205 cells), 99% of VCaP cells (n=10620 cells), 93% of PC-3 cells (n=6912 cells), and 100% of NCI-H1299 cells (n=19683). The assay revealed distinctive gene expression profiles for PBMCs, VCaP cells, PC-3 cells, and NCI-H1299 cells, as summarized in **Figure 2** and **Table 1**. Pairwise comparison revealed significant differences (q ≤ 0.05) in the expression of all markers between PBMCs and tumor cells, with the exception of SLFN11 and AR-V7 between PBMCs and PC-3 cells, and EPCAM, PSA, and PSMA between PBMCs and NCI-H1299 cells. Only for a small percentage it was not possible to differentiate between PBMCs and tumor cells using specific thresholds, since 1-8% of tumor cells expressed hematopoietic markers and 7% of PBMCs were positive for at least one epithelial or prostate-specific marker.

**Figure 2.**
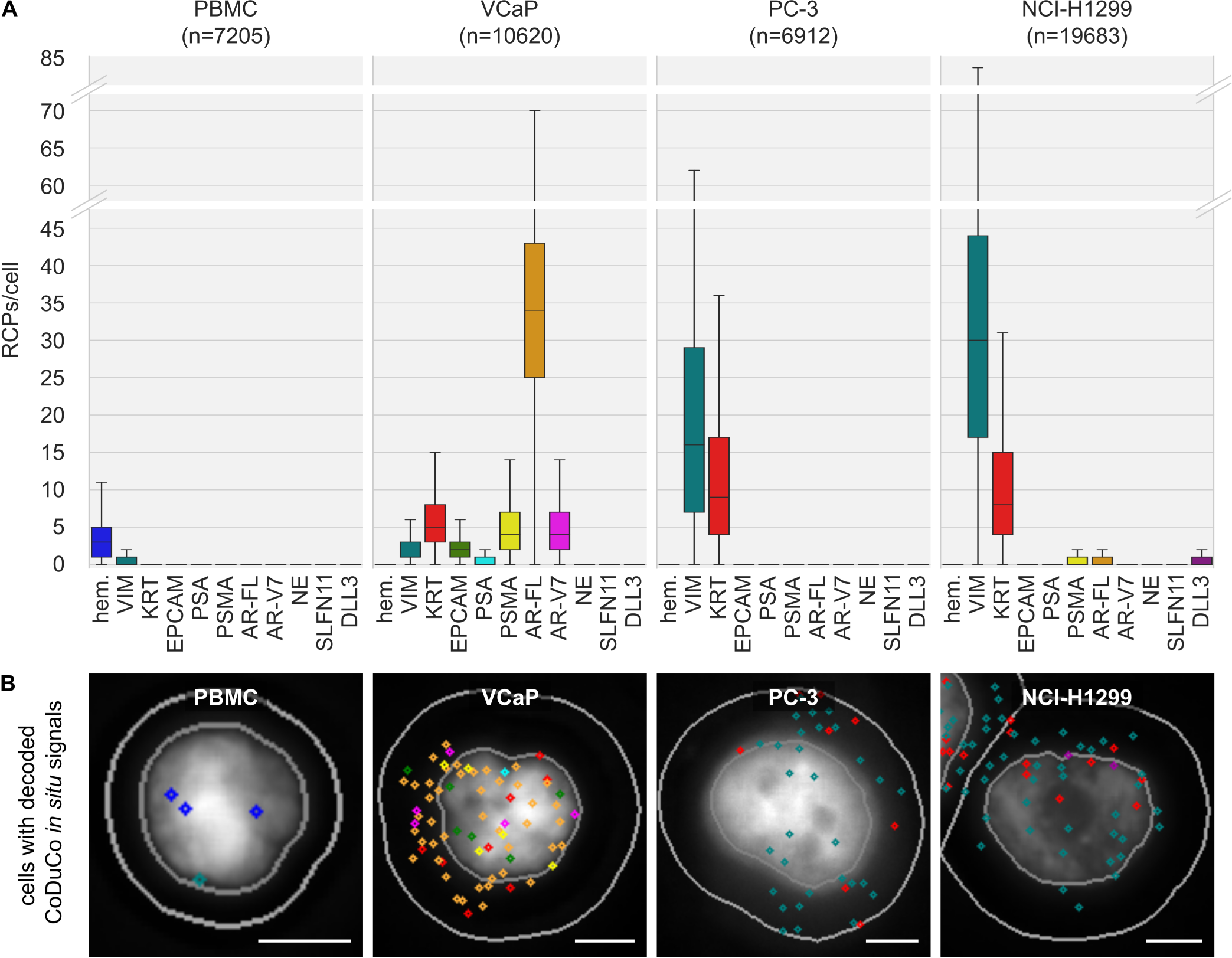
The novel CoDuCo in situ assay visualizes distinctive expression profiles in PBMCs, VCaP, PC-3, and NCI-H1299 cells. (A) The number of in situ signals per cell (RCPs/cell) is visualized for each marker and cell type/cell line. Outliers are not shown in the boxplots. (B) Exemplary images for each cell type, showing DAPI staining, nucleus and cell border outlines (gray), and decoded in situ signals as diamond shapes with the same color scheme as for the boxplots (e.g. blue diamond shapes represent hematopoietic markers). Scale bar 5 µm.

**Table 1.** CoDuCo in situ expression patterns in PBMCs and tumor cell lines VCaP, PC-3, and NCl-H1299.

| Marker | PBMC<br>(n = 7205) |  | VCaP<br>(n = 10620) |  | PC-3<br>(n = 6912) |  | NCI-H1299<br>(n = 19683) |  | q-value<br>PBMC versus |  |  |
| --- | --- | --- | --- | --- | --- | --- | --- | --- | --- | --- | --- |
|  | Positi-<br>vity<br>[%] | Median<br>(Q1, Q3)<br>[RCPs/cell] | Positi-<br>vity<br>[%] | Median<br>(Q1, Q3)<br>[RCPs/cell] | Positi-<br>vity<br>[%] | Median<br>(Q1, Q3)<br>[RCPs/cell] | Positi-<br>vity<br>[%] | Median<br>(Q1, Q3)<br>[RCPs/cell] | VCaP | PC-3 | NCI-H1299 |
| **hem.** | 86 | 3 (1-5) | 2 | 0 (0-0) | 8 | 0 (0-0) | 1 | 0 (0-0) | **0.000** | **0.000** | **0.000** |
| **VIM** | 32 | 0 (0-1) | 75 | 1 (1-3) | 92 | 16 (7-29) | 99 | 30 (17-44) | **0.000** | **0.000** | **0.000** |
| **KRT** | 4 | 0 (0-0) | 94 | 5 (3-8) | 90 | 9 (4-17) | 97 | 8 (4-15) | **0.000** | **0.000** | **0.000** |
| **EPCAM** | 0 | 0 (0-0) | 79 | 2 (1-3) | 21 | 0 (0-0) | 0 | 0 (0-0) | **0.000** | **0.000** | 0.848 |
| **PSA** | 0 | 0 (0-0) | 44 | 0 (0-1) | 3 | 0 (0-0) | 1 | 0 (0-0) | **0.000** | **0.000** | 0.501 |
| **PSMA** | 1 | 0 (0-0) | 93 | 4 (2-7) | 15 | 0 (0-0) | 28 | 0 (0-1) | **0.000** | **0.000** | **0.000** |
| **AR-FL** | 1 | 0 (0-0) | 99 | 34 (25-43) | 20 | 0 (0-0) | 32 | 0 (0-1) | **0.000** | **0.000** | **0.000** |
| **AR-V7** | 1 | 0 (0-0) | 90 | 4 (2-7) | 1 | 0 (0-0) | 1 | 0 (0-0) | **0.000** | 0.906 | 0.673 |
| **NE** | 4 | 0 (0-0) | 12 | 0 (0-0) | 3 | 0 (0-0) | 5 | 0 (0-0) | **0.000** | **0.003** | **0.014** |
| **SLFN11** | 7 | 0 (0-0) | 16 | 0 (0-0) | 6 | 0 (0-0) | 14 | 0 (0-0) | **0.000** | 0.415 | **0.000** |
| **DLL3** | 1 | 0 (0-0) | 14 | 0 (0-0) | 6 | 0 (0-0) | 26 | 0 (0-1) | **0.000** | **0.000** | **0.000** |
| **total** | 89 | 3 (2-6) | 99 | 55 (41-69) | 93 | 28 (15-48) | 100 | 41 (24-61) | **0.000** | **0.000** | **0.000** |

Positivity = percentage of cells with $\geq 1$ RCP/cell for the respective marker.
q-value from pairwise comparison between PBMCs and respective cancer cell lines using Dunn's Test with Benjamini/Hochberg correction. q-values <0.05 are highlighted in bold font.
hem.: pooled hematopoietic markers (PTPRC, ITGAM, FCGR3, CD4, ITGB2); KRT: pooled KRT markers (KRT8, KRT18, KRT19); NE: pooled neuroendocrine markers (SYP, CHGA, NCAM1).

In detail, pooled hematopoietic markers were expressed with an overall median of 3 RCPs/cell (IQR 1-5) and were detected in 86% of PBMCs. 2% of VCaP, 8% of PC-3, and 1% of NCI-H1299 cells were positive for pooled hematopoietic markers, but with very low overall expression levels (median 0 RCPs/cell, IQR 0-0). The assay detected low expression of epithelial and prostate-specific transcripts (KRT, EPCAM, PSA, PSMA, AR-FL, and AR-V7) in up to 4% of PBMCs, each (overall median 0 RCPs/cell, IQR 0-0). All epithelial and prostate-specific markers were expressed in VCaP cells. PSA had the lowest expression with a median of 0 RCPs/cell (IQR 0-1) and positivity in 44% of cells. AR-FL had the highest expression with a median of 34 RCPs/cell (IQR 25-43) and positivity in 99 % of cells. 90% of PC-3 cells expressed KRT, with a median overall expression of 9 RCPs/cell (IQR 4-17). Low expression of EPCAM and prostate-specific markers was detected in up to 21% of PC-3 cells, each (overall median 0 RCPs/cell, IQR 0-0). 97% of NCI-H1299 cells expressed KRT, with a median overall expression of 8 RCPs/cell (IQR 4-15). Low expression of PSA and AR-V7 was detected in 1% of NCI-H1299 cells, each (overall median 0 RCPs/cell, IQR 0-0). PSMA and AR-FL were detected in 28% and 32% of NCI-H1299 cells, respectively (overall median 0 RCPs/cell, IQR 0-1). VIM was detected in 32% of PBMCs, with a median overall expression of 0 RCPs/cell (IQR 0-1), in 75% of VCaP cells, with a median of 1 RCPs/cell (IQR 1-3), in 92% of PC-3 cells, with a median of 16 RCPs/cell (IQR 7-29), and in 99% of NCI-H1299 cells, with a median of 30 RCPs/cell (IQR 17-44). Low expression of pooled neuroendocrine markers, SLFN11, and DLL3, was detected in 4%, 7%, and 1% of PBMCs, 12%, 16%, and 14% of VCaP cells, and 3%, 6%, and 6% of PC-3 cells, respectively, with median overall expression of 0 RCPs/cell (IQR 0-0). Neuroendocrine markers and SLFN11 were detected in 5% and 14% of NCI-H1299 cells, respectively (overall median 0 RCPs/cell, IQR 0-0), and DLL3 in 26% of NCI-H1299 cells (overall median 0 RCPs/cell, IQR 0-1).

### Classifier training and evaluation

To create a ground truth for the classifier, a data set of 137871 cells was used, derived from 6 blood samples of healthy controls without (n=3) or with (n=3) spiked-in VCaP and PC-3 cells. Spiked blood samples were processed using the CytoGen Smart Biopsy Cell Isolator, transcripts were visualized and counted by CoDuCo *in situ* hybridization and CellProfiler image analysis, and all detected cells in the dataset were manually classified into the following 5 classes through expert evaluations: CTCs, PBMCs, artefacts, *in situ* false-positive cells, and *in situ* negative cells (**Figure 3**).

**Figure 3.**
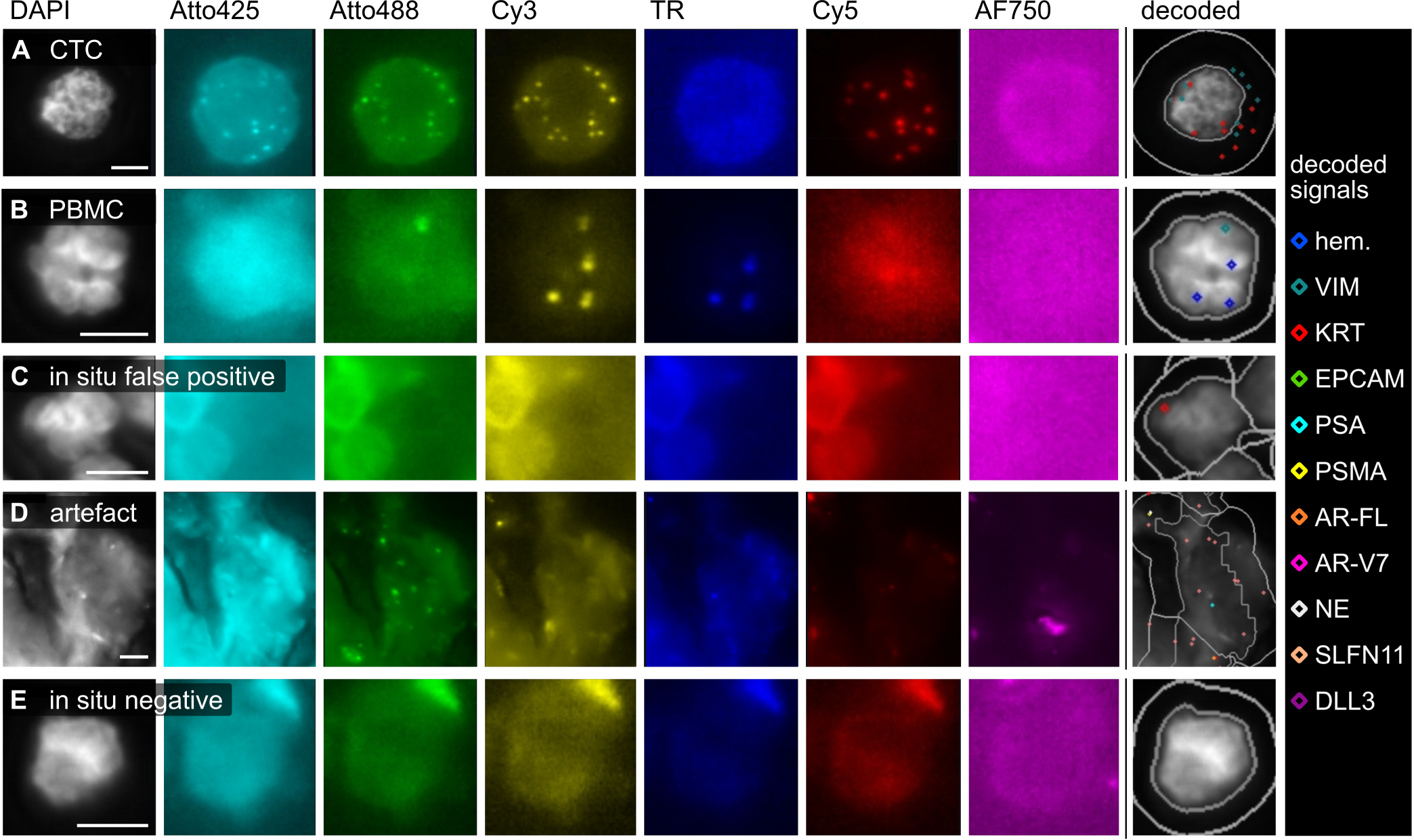
Based on the decoded in situ signals, the ground truth dataset was annotated. (A-E) For each of the 5 classes, one example is shown. (C) Objects with false-positive in situ signals (e.g. due to high autofluorescence) were classified as in situ false-positive (intact nuclear morphology) or (D) artefact (e.g. dust particles). (E) Objects without decoded in situ signals were classified as in situ negative. Scale bar 5 µm.

The ground truth dataset of 137871 cells was classified into the following classes: 673 CTCs (0.49%), 43219 PBMCs (31.35%), 156 artefacts (0.11%), 2514 *in situ* false-positive cells (1.82%), and 91309 *in situ* negative cells (66.23%). For sufficient representation of all classes in the training dataset, we randomly selected 1000 cells at a ratio of 2/5/1/1/1 (200 CTCs, 500 PBMCs, 100 of each of the remaining classes). After training a random forest classifier, we evaluated it in the remaining test dataset of 136871 cells. Confusion matrix and evaluation metrics are visualized in **Figure 4**. In the control samples, the classifier reached a high recall (0.89), precision (0.88), F1-score (0.89) and specificity (1.00) for CTCs.

**Figure 4.**
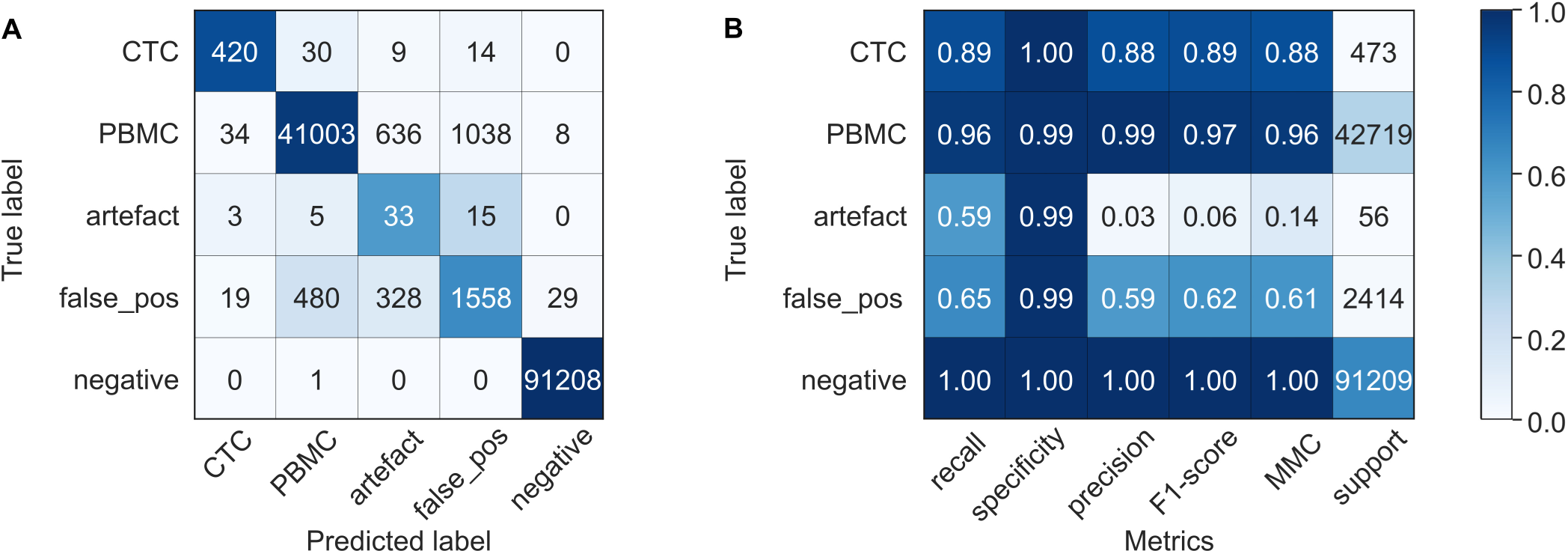
Classifier evaluation (test dataset). A random forest classifier was trained and tested on blood samples of healthy controls with and without spiked-in tumor cells. Results of the classifier evaluation in the test dataset are summarized in a confusion matrix (A) and classification report (B). In the confusion matrix, the colors indicate values normalized to the class support size (i.e. the number of actual occurrences in each group, as indicated in panel B), while the annotation shows non-normalized values. false_pos = cells without true-positive in situ signals; negative = cells with no in situ signals detected by CellProfiler analysis; recall = fraction of correctly identified positives; specificity = fraction of correctly identified negatives; precision = accuracy of positive predictions; F1-score = harmonic mean of precision and recall; MCC = Matthews Correlation Coefficient.

### The classifier identified patient CTCs with a recall of 0.76 and specificity of 0.99

Finally, we tested the complete workflow, including blood collection, CTC enrichment, *in situ* hybridization, image analysis, cell classification, and expert revision, on three patient samples (PC-13, PC-15, PC-16). In total, 17756 cells were detected by automated image analysis. During expert revision, 49 of them were identified as CTCs, of which 37 (76%) were also recognized by the classifier, resulting in a recall of 0.76 as visualized in **Figure 5**. Among 17707 non-CTCs, 177 false-positive CTCs were reported by the classifier, corresponding to a specificity of 0.99. With 37 true-positive and 177 false-positive CTCs, the classifier reached a precision of 0.17.

**Figure 5.**
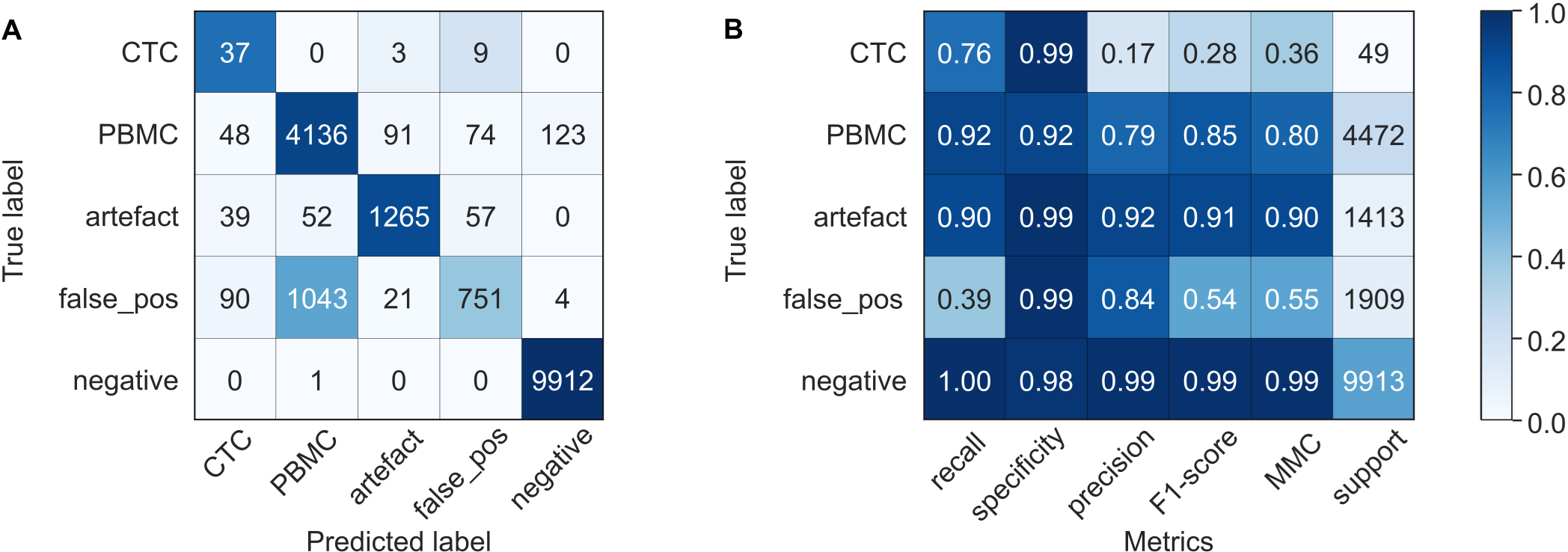
Classifier evaluation on three patient samples. A random forest classifier was trained on blood samples of healthy controls with and without spiked-in tumor cells. Results of the classifier evaluation on three patient samples are summarized in a confusion matrix (A) and classification report (B). In the confusion matrix, the colors indicate values normalized to the class support size (i.e. the number of actual occurrences in each group, as indicated in panel B), while the annotation shows non-normalized values. false_pos = cells without true-positive in situ signals; negative = cells with no in situ signals detected by CellProfiler analysis; recall = fraction of correctly identified positives; specificity = fraction of correctly identified negatives; precision = accuracy of positive predictions; F1-score = harmonic mean of precision and recall; MCC = Matthews Correlation Coefficient.

Comparative analysis of expression patterns between false-negative (24%) and true-positive (76%) CTCs revealed major differences (**Table 2**). In CTCs that were missed by the classifier, the total number of automatically detected *in situ* signals per cell was significantly decreased, with a median of 4 RCPs/cell (IQR 3-7) compared to a median of 35 RCPs/cell (IQR 18-67) in true-positive CTCs (q ≤ 0.0001). Furthermore, there was a significant decrease in expression levels of KRT (q ≤ 0.0001), PSA (q ≤ 0.001), AR-FL (q ≤ 0.001), and AR-V7 (q ≤ 0.05) in false-negative CTCs. 75% of false-negative CTCs showed no KRT expression, while KRT was expressed in 97% of true-positive CTCs. In KRT-negative CTCs, the classifier performance was significantly decreased, with a recall of 0.10 compared to 0.92 for KRT-positive CTCs.

**Table 2.**
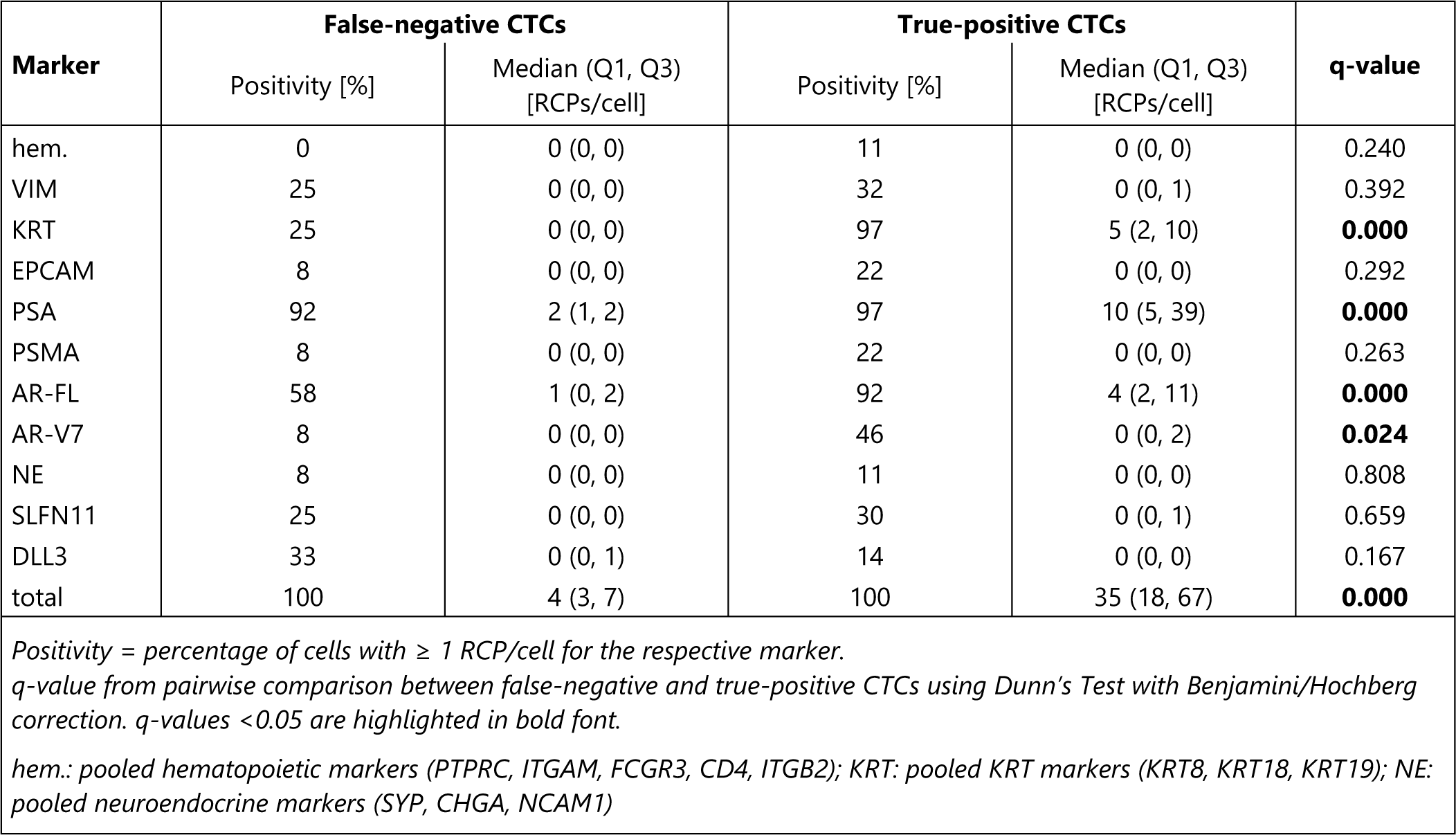
Comparison of expression patterns in patient CTCs that were missed or recognized by the classifier.

### CoDuCo revealed interpatient CTC heterogeneity and captured neuroendocrine CTCs and CTC-clusters

Interpatient heterogeneity was observed regarding CTC count, presence of CTC clusters, as well as expression patterns. Exemplary images of patient CTCs are depicted in **Figure 6**. We found 8 CTCs in sample PC-13, 3 CTCs in PC-15, and 38 CTCs in PC-16. In PC-16, most CTCs (26 of 38) were found in clusters of up to 9 CTCs (**Figure 6 D**), while no CTC-clusters were found in the other samples.

**Figure 6.**
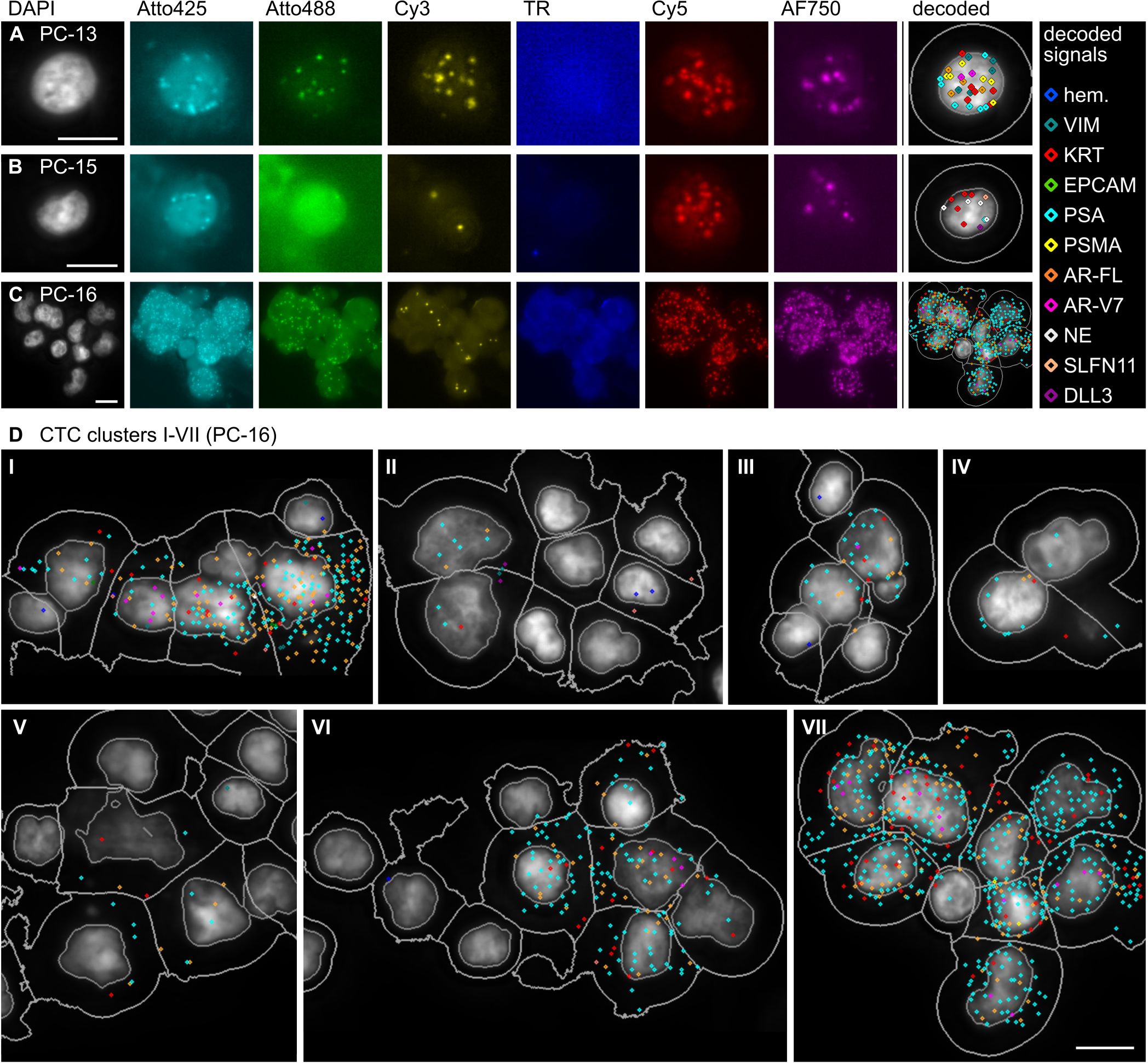
Exemplary images of patient CTCs. (A-C) Normalized pseudocolored fluorescence images and overlay images showing the DAPI channel with outlines of nuclei, cell borders, and decoded in situ signals as detected by CellProfiler analysis. (D) Overlay images with decoded in situ signals in seven CTC-clusters that were found in PC-16. Only cluster IV and VII consisted exclusively of CTCs. The other clusters were mixed and contained PBMCs and/or in situ negative cells as well as CTCs. Scale bar 10 µm.

The interpatient differences in CTC numbers and their expression level are summarized in **Table 3** and **Figure 9 C**. PC-13 CTCs were characterized by high VIM and KRT expression and medium to low expression of prostate-specific markers, such as PSMA and AR-V7. PC-16 CTCs showed high expression of PSA, medium expression of AR-FL, low expression of KRT, and very low overall expression of VIM, PSMA, and AR-V7. In contrast, CTCs in PC-15 expressed no prostate-specific markers but expressed KRT and neuroendocrine markers at a high level. Pairwise comparison between patient samples revealed that VIM and PSMA expression was significantly increased in PC-13 (q ≤ 0.05), PSA expression was significantly increased in PC-16, and the expression of pooled neuroendocrine markers, SLFN11, and DLL3 was significantly increased in PC-15. No significant difference in KRT expression was detected. In detail, the median number of RCPs/cell in CTCs of PC-13 was 8 for VIM (IQR 4-15), 8 for KRT (IQR 2-14), 2 for PSA (IQR 0-8), 4 for PSMA (IQR 2-5), 2 for AR-FL (IQR 1-3), and 0 for AR-V7 (IQR 0-1). In CTCs of PC-15, the median number of RCPs/cell was 7 for KRT (IQR 6-8), 0 for EPCAM (IQR 0-1), 7 for pooled neuroendocrine markers (IQR 6-10), 2 for SLFN11 (IQR 1-3), and 1 for DLL3 (IQR 0-2). In CTCs of PC-16, the median number of RCPs/cell was 2 for KRT (IQR 1-5), 10 for PSA (IQR 4-38), 4 for AR-FL (IQR 1-10), and 0 for AR-V7 (IQR 0-1).

**Table 3.** Interpatient comparison of CTC expression patterns.

| Marker | PC-13 CTCs (n = 8) |  | PC-15 CTCs (n = 3) |  | PC-16 CTCs (n = 38) |  | q-values |  |  |
| --- | --- | --- | --- | --- | --- | --- | --- | --- | --- |
|  | Positivity [%] | Median (Q1, Q3) [RCPs/cell] | Positivity [%] | Median (Q1, Q3) [RCPs/cell] | Positivity [%] | Median (Q1, Q3) [RCPs/cell] | PC-13 & PC-15 | PC-13 & PC-16 | PC-15 & PC-16 |
| hem. | 0 | 0 (0-0) | 0 | 0 (0-0) | 3 | 0 (0-0) | 1.000 | 1.000 | 1.000 |
| VIM | 88 | 8 (4-15) | 33 | 0 (0-0) | 3 | 0 (0-0) | **0.031** | **0.000** | 0.268 |
| KRT | 75 | 8 (2-14) | 100 | 7 (6-8) | 76 | 2 (1-5) | 0.628 | 0.195 | 0.195 |
| EPCAM | 25 | 0 (0-0) | 33 | 0 (0-1) | 8 | 0 (0-0) | 0.591 | 0.319 | 0.319 |
| PSA | 62 | 2 (0-8) | 0 | 0 (0-0) | 100 | 10 (4-38) | 0.214 | **0.026** | **0.010** |
| PSMA | 88 | 4 (2-5) | 0 | 0 (0-0) | 5 | 0 (0-0) | **0.001** | **0.000** | 0.845 |
| AR-FL | 75 | 2 (1-3) | 0 | 0 (0-0) | 89 | 4 (1-10) | 0.158 | 0.158 | **0.023** |
| AR-V7 | 50 | 0 (0-1) | 0 | 0 (0-0) | 32 | 0 (0-1) | 0.394 | 0.506 | 0.394 |
| NE | 0 | 0 (0-0) | 100 | 7 (6-10) | 0 | 0 (0-0) | **0.000** | 1.000 | **0.000** |
| SLFN11 | 25 | 0 (0-0) | 67 | 2 (1-3) | 3 | 0 (0-0) | **0.036** | 0.072 | **0.001** |
| DLL3 | 12 | 0 (0-0) | 67 | 1 (0-2) | 0 | 0 (0-0) | **0.001** | 0.194 | **0.000** |
| **total** | 100 | 39 (21-50) | 100 | 20 (16-22) | 100 | 16 (7-54) | 0.955 | 0.955 | 0.971 |

Positivity = percentage of cells with $\geq 1$ RCP/cell for the respective marker.
q-value from pairwise interpatient comparison using Dunn's Test with Benjamini/Hochberg correction. q-values <0.05 are highlighted in bold font.
hem.: pooled hematopoietic markers (PTPRC, ITGAM, FCGR3, CD4, ITGB2); KRT: pooled KRT markers (KRT8, KRT18, KRT19); NE: pooled neuroendocrine markers (SYP, CHGA, NCAM1)

### CoDuCo *in situ* revealed intrapatient CTC heterogeneity

On a single cell level, individual CTCs showed very heterogeneous expression patterns as visualized in the clustermaps and CTC images in **Figure 7** and **Figure 8**. In PC-13, multiple CTCs had remarkably high coexpression of VIM and KRT in the absence of AR-V7, while others had more prominent expression of prostate-specific markers, especially PSA and AR-V7 (**Figure 7 A**). The three CTCs that were detected in patient sample PC-15 all showed coexpression of KRT and neuroendocrine markers (**Figure 7 B**). In PC-16, all CTCs expressed PSA, but the expression level ranged from 1 RCP/cell to 80 RCPs/cell. Similarly, most CTCs were positive for KRT and AR-FL, with expression levels ranging from 0-20 RCPs/cell and 0-67 RCPs/cell, respectively. Furthermore, 12/38 CTCs expressed AR-V7. While the median overall expression of AR-V7 was 0 RCPs/cell (IQR 0-1), in the subset of AR-V7-positive CTCs, the median expression was 3 RCPs/cell (IQR 2-4) (**Figure 8**).

**Figure 7.**
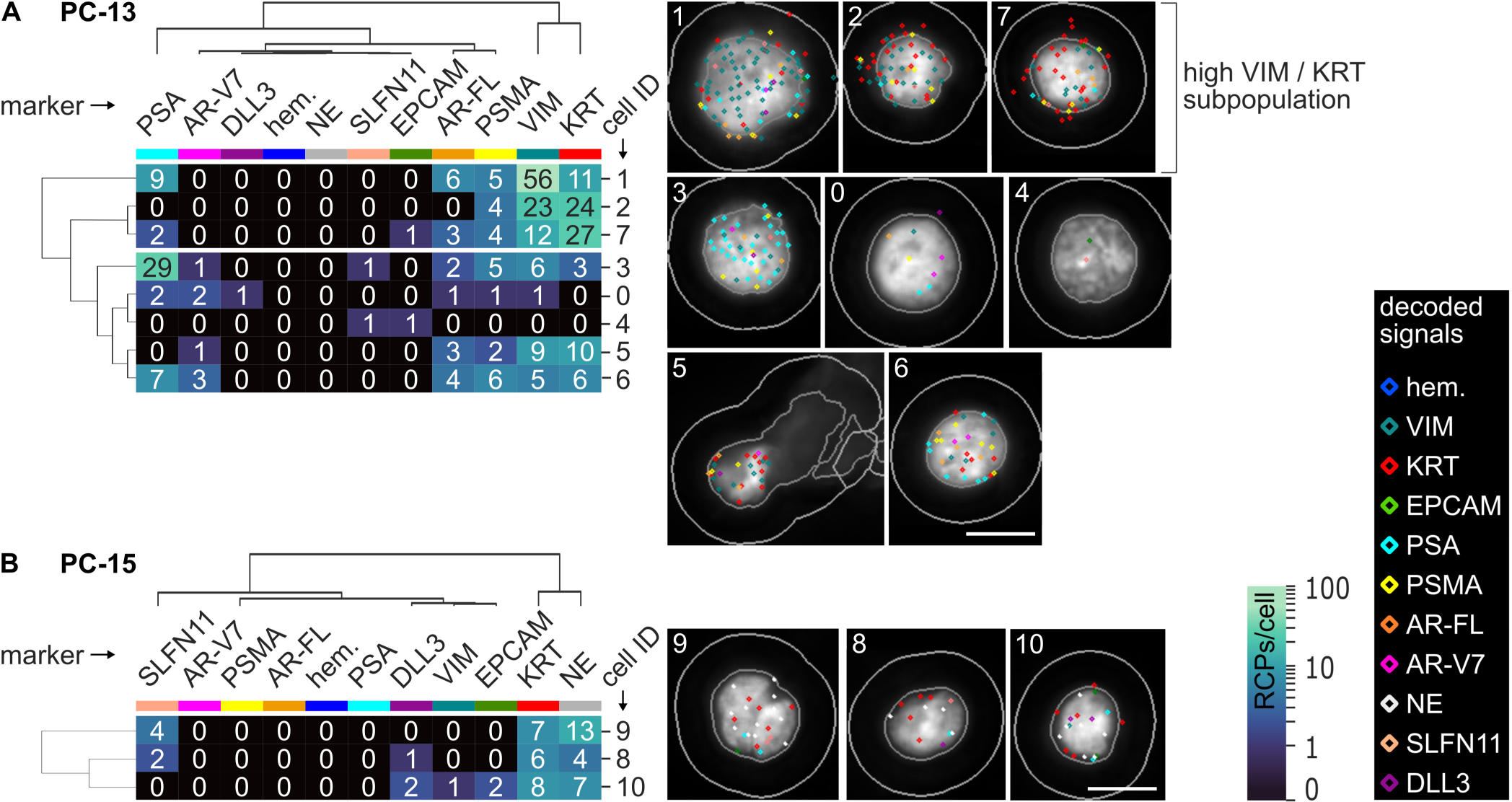
All detected CTCs of patient PC-13 (A) and PC-15 (B). Dendrograms as well as the order of cells (rows) and markers (columns) are based on hierarchical clustering of CTCs. The number of in situ signals/cell is visualized in the heatmaps and corresponding CTC thumbnails. The thumbnails show the DAPI image with cell ID number, outlines of nuclei, cell borders, and decoded in situ signals as detected by CellProfiler analysis. The colors of decoded in situ signals is indicated for each marker on top of the clustermap. For instance, red dots are KRT transcripts. We found heterogeneous subpopulations of CTCs (high KRT/VIM and PSA/AR-V7, respectively) in patient PC-13 (A) and neuroendocrine CTCs in PC-15 (B). Pools of 3-5 genes were used for hematopoietic hem. (PTPRC, ITGAM, FCGR3A&B, CD4, ITGB2), KRT (KRT8, KRT18, KRT19), and neuroendocrine NE markers (SYP, CHGA, NCAM1). Scale bar 10 µm.

**Figure 8.**
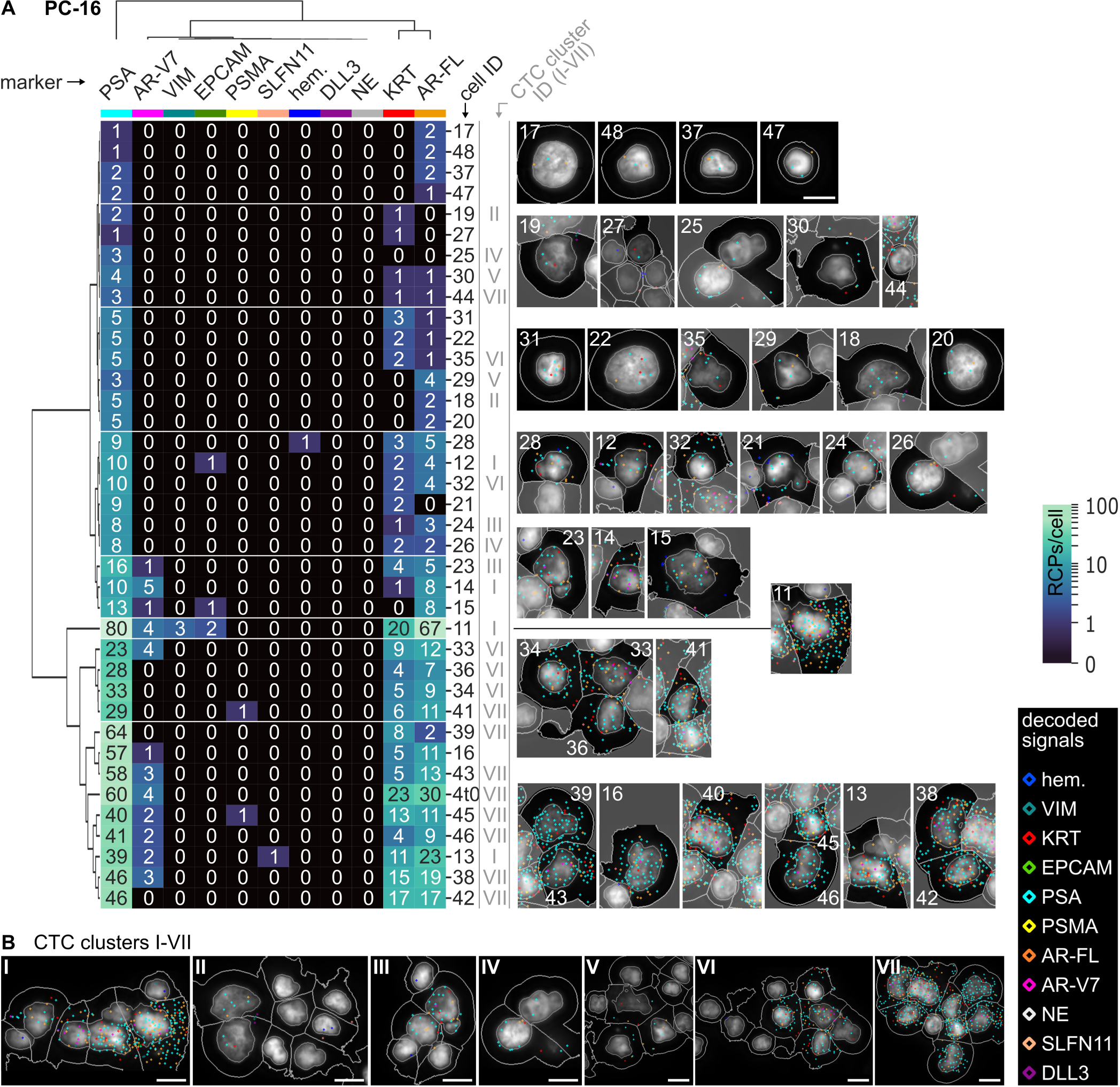
CTCs of patient PC-16. (A) Dendrograms as well as the order of cells (rows) and markers (columns) are based on hierarchical clustering of CTCs. The number of in situ signals/cell is visualized in the heatmaps and corresponding CTC thumbnails. The thumbnails show the DAPI image with cell ID number, outlines of nuclei, cell borders, and decoded in situ signals as detected by CellProfiler analysis. The colors of decoded in situ signals is indicated for each marker on top of the clustermap. (B) Overview of 7 CTC-clusters (IDs I-VII) detected in PC-16. Pools of 3-5 genes were used for hematopoietic hem. (PTPRC, ITGAM, FCGR3A&B, CD4, ITGB2), KRT (KRT8, KRT18, KRT19), and neuroendocrine NE markers (SYP, CHGA, NCAM1). Scale bar 10 µm.

### CoDuCo *in situ* showed high concordance with clinical parameters and AdnaTest

The results of the *in situ* assay were in line with clinical parameters (**Figure 9 A and C**). Total PSA measurements of 6.75 ng/ml in PC-13 and 904.06 ng/ml in PC-16 were also reflected by CTC numbers and PSA expression levels determined by *in situ* analysis, with PSA positivity in 5 of 8 CTCs and a median overall expression of 2 PSA RCPs/cell (IQR 0-8) in PC-13, compared to PSA positivity in all 38 CTCs and a median expression of 10 PSA RCPs/cell (IQR 4-38) in PC-16. In PC-15, a patient with treatment emergent small-cell neuroendocrine PC, CTCs expressing neuroendocrine markers (SYP, CHGA, NCAM1) and DLL3 co-occurred with elevated blood levels of neuron-specific enolase (1451 ng/ml) and chromogranin A (2764 ng/ml). Similarly, *in situ* CTC analysis showed high agreement with the AdnaTest results (**Figure 9 B and C**). PSA, PSMA, AR-FL/AR, and AR-V7 were detected in patient PC-13 and PC-16 by both assays. In patient PC-15, the assays were concordant regarding the absence of PSA, AR-FL/AR, and AR-V7. Interestingly, low-level PSMA expression was detected by the AdnaTest but not by *in situ* analysis.

**Figure 9.**
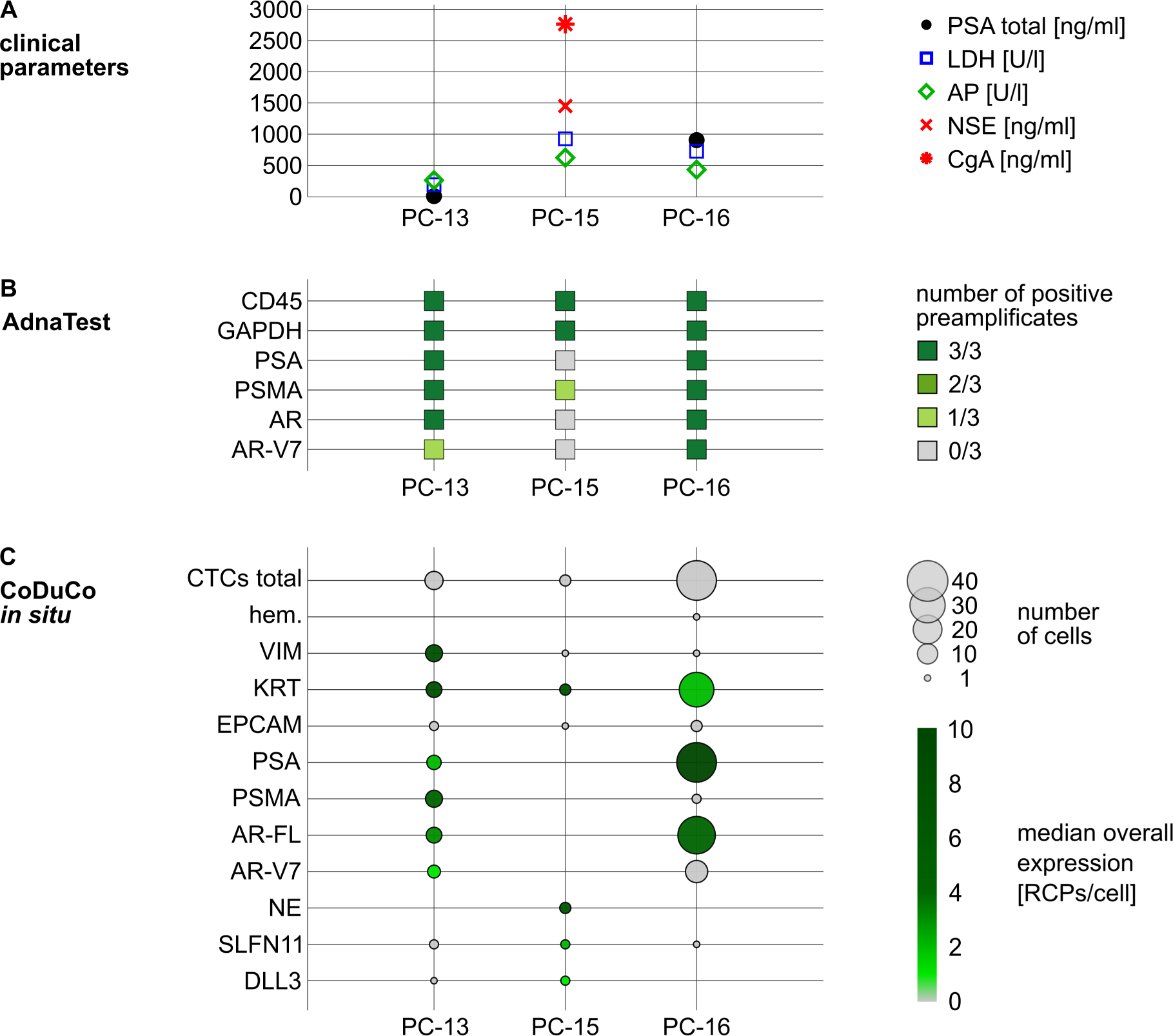
Visualization of clinical parameters, results of CTC-analyses by AdnaTest and CoDuCo in situ assay for patients PC-13, PC-15, and PC-16. (A) Blood levels of total PSA [ng/ml], lactate dehydrogenase (LDH) [U/l], alkaline phosphatase (AP) [U/l], neuron-specific enolase (NSE) [ng/ml], and chromogranin A (CgA) [ng/ml], where available. (B) AdnaTest results. For each sample, 3 preamplifications of isolated and reverse-transcribed RNA were performed for full coverage. The absence or presence of PSA, PSMA, AR (=AR-FL), and AR-V7 transcripts is visualized by colored squares (dark green = detected in all preamplificates; gray = not detected). (C) Bubble plot visualizing the number (bubble size) and median overall expression levels (color scale) of in situ analyzed CTCs (dark green = high expresses; gray = low expressed). hem.: pooled hematopoietic markers (PTPRC, ITGAM, FCGR3, CD4, ITGB2); KRT: pooled KRT markers (KRT8, KRT18, KRT19); NE: pooled neuroendocrine markers (SYP, CHGA, NCAM1).

## DISCUSSION

Our data show that CTCs can be identified and characterized by our novel CoDuCo *in situ* assay, which targets up to 11 markers in a multiplex fashion. Multiple predictive biomarkers are detected simultaneously, which previously was not feasible. Of utmost importance is the assay’s capability to detect neuroendocrine markers, as this remains challenging with conventional antibody staining procedures. Quantitative assessment of expression levels of CTCs is possible, and the machine learning classifier is a powerful tool to recognize CTCs. The assay provides single cell resolution RNA expression data and images of cell morphology, which facilitates the identification of single CTCs and CTC clusters and strikingly reveals intrapatient heterogeneity.

### Neuroendocrine transdifferentiation is detectable in CTCs

The increased multiplex capacity of the CoDuCo *in situ* approach enabled us to visualize in total 11 markers, either as single transcripts (e.g. SLFN11, DLL3), or pooled markers (e.g. SYP, CHGA, NCAM1). The combination of markers increases the informative value which can be obtained from CTCs. Simultaneous assessment of 11 targets is difficult to accomplish with conventional antibody staining or would require sophisticated and expensive targeted proteomics mass cytometry techniques (46). Using our CoDuCo assay, we could clearly identify CTCs, but more importantly, we could investigate multiple resistance markers on a single cell level. Of high clinical importance is the exploration of neuroendocrine transdifferentiation, which is an AR-independent resistance mechanism in CRPC. While the incidence of neuroendocrine PC is rising, its early detection and treatment remain difficult and improvements are urgently needed (8, 47, 48). Thanks to the CoDuCo *in situ* assay’s improved multiplexing capability, we were able to add markers relevant for neuroendocrine PC to our panel. We visualized SYP, CHGA, and NCAM1 transcripts as pooled neuroendocrine markers to identify CTCs with neuroendocrine features, as well as DLL3 and SLFN11 as potential predictive markers. Indeed, we successfully identified neuroendocrine CTCs in a patient with diagnosed treatment-emergent neuroendocrine PC (PC-15) in our proof of principle study. A large patient cohort would be needed to determine whether *in situ* CTC analysis can reveal neuroendocrine transdifferentiation earlier or with higher specificity than the serum markers currently used in the clinic. However, a clear advantage of our approach is the parallel detection of DLL3 and SLFN11 expression in neuroendocrine CTCs. SLFN11 expression is a potential positive predictive biomarker for platinum-based chemotherapy and PARP inhibitors, and thus relevant for neuroendocrine PC (49–51). In contrast, DLL3 has been associated with resistance against platinum-based chemotherapy (52, 53). Notably, DLL3 itself is a potential therapeutic target, which is currently investigated in several preclinical and clinical studies (16, 54). So far, only a limited number of studies investigated and confirmed the detection of DLL3 expression in CTCs of CRPC and small cell lung cancer patients (54–56). Although DLL3 expression in CTCs is representative of expression in matched metastatic tissue biopsies (54), conflicting data exists regarding DLL3’s predictive value in the context of DLL3-targeted therapy and further investigation is needed (56). In the case presented here, we detected three neuroendocrine CTCs. SLFN11 and DLL3 were detected in two of three CTCs, respectively. Coexpression of SLFN11 and DLL3 was observed in one CTC, while the remaining CTCs expressed either SLFN11 or DLL3. An interesting finding is that neuroendocrine CTCs still expressed keratin, despite having no prostate-specific transcripts such as PSA, PSMA or AR-FL. This might be important for other CTC-based assays that use keratin staining as inclusion marker, implying they can still identify CTCs in neuroendocrine PC patients. Although these data of a single case merely confirm the capability to detect neuroendocrine markers in CTCs, the findings suggest that this assay may bring novel insights regarding the treatment options and resistance mechanisms in neuroendocrine PC.

### PSMA expression is detectable in CTCs

Similarly, the increased multiplex capacity of the CoDuCo approach enabled us to include PSMA as additional prostate-specific marker. Novel PSMA-targeting drugs, such as the radioligand agent Lutetium-177 PSMA-617 (trade name: Pluvicto) (18), are now available, and others are under investigation (17, 57). Predictive biomarkers are urgently needed, as the expression of PSMA is highly heterogeneous and dynamic (58). PSMA upregulation by ADT, by inhibition of the PI3K/Akt/mTOR pathway, and by cell stress through DNA damaging treatments or DNA damage response inhibitors have been described, which implies potential benefits of combination treatments (59). Thus, visualizing PSMA expression in CTCs might not only prove useful for patient stratification, but may, in the context of a comprehensive multi-analyte liquid biopsy approach, lead to a better understanding of PSMA regulation and potential implications for combination treatments (60). In the data presented here, we detected PSMA-expressing CTCs in two patients but none in the patient with treatment-emergent neuroendocrine PC, where we detected three CTCs expressing KRT and neuroendocrine markers, but no prostate-specific markers. These findings are in line with published data demonstrating decreased or absent PSMA expression in AR-negative PC (59, 61). In contrast to the CoDuCo *in situ* results, PSMA expression was detected by the AdnaTest assay in all samples. The discordance between the two assays may point to differences in sensitivity and/or specificity and should be investigated in a larger cohort of PC patients and healthy controls. Importantly, in high risk/advanced PC patients the current gold standard technique for PSMA detection is a whole-body imaging with positron emission tomography (PET) using small amounts of radioactive tracers, such as 68Ga-PSMA-11 (62). Utilizing a liquid biopsy assay as an alternative to PET imaging with radioactive tracers would be highly beneficial for patients and healthcare providers. PSMA expressing CTCs might be used for patient stratification, monitoring the efficacy of Lutetium-177 PSMA-617 radioligand treatment, and identifying potential resistance mechanisms to this novel treatment option.

### AR-FL, AR-V7, and PSA are detectable and align with clinical serum markers

We designed our CoDuCo *in situ* assay to visualize the expression of AR-FL, AR-V7, and PSA, among others. In comparison to our previous study (23), the detection of AR-FL and PSA was optimized by increasing the number of PLPs per transcript from a single PLP to seven and six PLPs, respectively. As inhibition of the AR pathway is the mainstay of systemic PC treatment and AR-V7 is a well described resistance marker in up to 25% of CRPC patients, assessing patients’ AR status over the course of the treatment is highly relevant (63). PSA is an AR-regulated gene, meaning that its expression can be interpreted as a measurement for AR-pathway activity when monitoring the response to AR-targeted treatments (64). Furthermore, serum PSA is the most popular biomarker in PC and is routinely monitored in the clinical setting. We found high concordance between serum PSA levels and CoDuCo *in situ* derived expression levels of PSA-positive CTCs. This concordance supports our CoDuCo *in situ* results with actual clinical data and strengthens our assay.

### Single cell resolution reveals tumor heterogeneity in CTCs and CTC clusters

The CoDuCo *in situ* assay delivers single cell resolution RNA expression data and images of cell morphology, thereby enabling the identification of single CTCs and CTC clusters, and the exploration of tumor heterogeneity. We observed intrapatient CTC heterogeneity in two patients, namely, PC-13 and PC-16. In PC-16 we found substantial heterogeneity of KRT, PSA, AR-FL, and AR-V7 expression levels between CTCs. However, the expression levels of all these markers followed a similar gradient, from low expression of all markers to high expression of all markers, suggesting that the heterogeneity may have been caused by transcriptional activity overall, rather than multiple clones preexisting in the tumor mass (cellular plasticity). In contrast, PC-13 did not follow this pattern. Instead, there were at least two CTC cell populations, one with remarkably high coexpression of KRT and VIM in the absence of AR-V7, and one with more prominent expression of prostate-specific markers, especially PSA and AR-V7. This might indicate clonal heterogeneity of the tumor mass as described previously (65). Overall, CTC heterogeneity has been described as predictive biomarker in metastatic CRPC (66), highlighting the importance of single cell-based approaches like our CoDuCo *in situ* assay, which can visualize and quantify heterogeneity among CTCs. In PC-16, we detected CTC clusters with and without associated PBMCs or *in situ*-negative cells. The CTC clusters differed both in their expression patterns as well as their size, expressing KRT, PSA, AR-FL, and in some cases AR-V7, but no VIM, and the largest cluster contained nine CTCs. The detection and analysis of CTC clusters is highly relevant, as they are associated with high metastatic potential and poor outcomes (67–69) and drugs that target and disaggregate CTC clusters are under investigation in preclinical and clinical (NCT03928210) studies (70, 71).

### Image analysis and machine learning-based classification

A challenge for the CoDuCo assay is the complexity of image analysis. In our previous study, a four-channel fluorescence microscope limited our conventional *in situ* PLP hybridization approach to detecting only three markers, but image analysis was comparably simple (23). In our novel CoDuCo *in situ* assay, we used a seven-channel fluorescence microscope and dual-color combinations to detect *in situ* signals. In theory, this combinatorial approach can distinguish *in situ* signals with up to 15 unique color codes (72). However, with CoDuCo staining, a highly optimized, crosstalk-free filter setup is required to ensure specificity. As we observed some bleed-through of Cy5-labelled signals into the TexasRed channel, we used TexasRed only for a single color-code instead of five, ensuring high specificity.

To ease image analysis and sample evaluation, we developed a semi-automated image analysis pipeline using Python libraries and CellProfiler and established machine learning-assisted classification of cells. Using this pipeline, we decoded colocalized *in situ* signals and assigned them to cells that we detected based on their DAPI-stained nuclei. Although the automation of image analysis ensured the feasibility of the CoDuCo *in situ* analysis of CTCs, we still encountered technical limitations and therefore further optimizations will be indispensable. These limitations included wide variations of the intensity values of DAPI staining, *in situ* signals, but also background autofluorescence of patient samples. Although our CellProfiler pipeline used automated thresholding algorithms to detect nuclei and *in situ* signals, some parameters needed to be checked and adjusted for each individual sample to ensure optimal segmentation results. In addition to that, we stripped the *in situ* signals from the samples and created a background scan to be subtracted from the original images to increase the signal to background ratio (32). Both the manual adjustments and the background scan are time-consuming and therefore, optimizations would be worthwhile. The use of machine learning and deep learning-based tools for deconvolution, spot enhancement, and image segmentation (e.g. Cellpose, StarDist, ilastik, DeepSpot, Deconwolf) might improve the detection of nuclei and *in situ* signals and speed up the analysis workflow (73–77). Moreover, as our CoDuCo *in situ* assay involved no cytoplasm or membrane staining, we enlarged detected nuclei, based on their size, by a specified number of pixels to capture the whole cells. Implementing membrane or cytoplasmic staining, such as cell painting tools, to guide the detection of cell borders, would be another potential improvement to the image analysis workflow (78, 79).

To train a classifier for CTC detection by supervised machine learning, we first had to create a ground-truth dataset. We used blood samples of healthy controls with and without spiked-in VCaP and PC-3 cells, enriched them for CTCs using CytoGen’s Smart Biopsy Cell Isolator, and performed CoDuCo *in situ* analysis. We manually annotated the dataset and trained a random forest classifier using the CellProfiler Analyst software. When tested on three samples of PC patients, the classifier reached a recall of 0.76 and specificity of 0.99, meaning that 76% of CTCs and 99% of non-CTCs were correctly recognized, and a precision of 0.17, meaning that 17% of predicted CTCs were true CTCs. The observed discrepancy of performance metrics between the healthy controls’ spike-in samples and the patient samples indicated that training and test dataset were not representative for patient samples. This finding was expected as there is a large difference between cultured cancer cell lines and patient CTCs (80). The importance of a representative training dataset to correctly identify CTCs was also noted by others, who used deep-learning convolutional neural networks to classify CTCs, either via deep-learning or by operator reviewed CTCs (81). Since the identification of CTCs can be difficult even for experts, the creation of a ground truth dataset is very challenging. Therefore, including additional samples of healthy controls is of particular importance to minimize false-positive CTC calls and improve classifier performance overall (82). Also, identification of CTCs might be improved by including morphological features, so that classification is based on molecular (mRNA *in situ* signals) and cellular (e.g. nucleus shape) features. Eventually re-training of the classifier is needed, using a more representative dataset (82). The most suitable ground truth dataset for classifier training will be a collection of several hundred CoDuCo *in situ* hybridized samples from healthy controls and PC patients with a large number of expert-reviewed patient CTCs.

### Conclusion

We demonstrated the feasibility of a novel CoDuCo *in situ* approach which identifies CTCs with high sensitivity and specificity. CoDuCo staining increases the multiplex capacity of this assay, allowing us to visualize a more comprehensive panel of transcripts, including neuroendocrine, epithelial, prostate-specific, mesenchymal, and hematopoietic markers. The transcripts were selected to inform about diverse resistance mechanisms (AR-V7 expression, neuroendocrine transdifferentiation), druggable targets and predictive markers (PSMA, DLL3, SLFN11), and cancer-related processes such as epithelial mesenchymal transition (VIM, KRT). To ensure practical applicability, we implemented semi-automated image analysis combined with machine learning-assisted CTC classification. A unique advantage of the CoDuCo *in situ* assay is the combination of high multiplex capacity and microscopy-based single-cell analysis, which is instrumental to simultaneously identify and characterize CTCs, detect CTC clusters, and visualize CTC heterogeneity. Ultimately, the assay is a promising tool for tracking the evolving molecular alterations linked to drug response and resistance in PC.

## ACKNOWLEDGEMENTS

The data presented in this study were also a component of the dissertation submitted by Lilli Bonstingl to Medical University of Graz in 2023, entitled ‘Tracking the resistance: liquid biopsy monitoring of drug resistance in metastatic prostate cancer’. The authors wish to formally acknowledge the contributions of Peter Abuja, and Kurt Zatloukal, and the technical support by Daniel Kummer, Laurin Herbsthofer, and Michael Gruber, their guidance and assistance was instrumental in the research and preparation of this manuscript. The authors thank Gerlinde Gornicec, Carina Kreuter, Sylvia Tripolt, Karin Groller, and Lisa Jaritz from the study coordination team of the Division of Oncology, Medical University of Graz, for their excellent support in patient enrolment. We acknowledge the use of ChatGPT-4, provided by OpenAI, only for enhancing the writing in specific sections of this manuscript. Figures were created with BioRender.com and Inkscape.

## FUNDING

This work was performed within the K1 COMET Competence Center CBmed, which is funded by the Federal Ministry of Transport, Innovation and Technology (BMVIT); the Federal Ministry of Science, Research and Economy (BMWFW), Land Steiermark (Department 12, Business and Innovation), the Styrian Business Promotion Agency (SFG), and the Vienna Business Agency. The COMET program is executed by the Austrian Research Promotion Agency (FFG). Authors were supported by CBmed and the Medical University of Graz via the PhD program Advanced Medical Biomarker Research (AMBRA) and the Doctoral School in Translational Molecular and Cellular Biosciences. The research was supported by a grant of the Verein für Krebskranke of the Medical University of Graz and by MEFO Graz, the Medical research Funding Society of the Medical University of Graz (Austria).

## CONFLICT OF INTEREST

The authors declare that there are no conflicts of interest.

## Notes

### Competing Interest Statement

The authors have declared no competing interest.

